# The *SAUERKRAUT* transposable element accelerates Arabidopsis floral transition

**DOI:** 10.64898/2026.04.03.715599

**Authors:** Joram A. Dongus, Yu Him Tang, Annabel D. van Driel, Michael A. Schon, Eslin T. Pleunis, Kilian Duijts, Joy Debnath, Iko T. Koevoets, Pinelopi Kokkinopoulou, Thijs de Zeeuw, Sofía Ortega, A. Jessica Meyer, Anne M. Blok, René Boesten, Christa Testerink

**Author notes:** Plant Stress Resilience, Institute of Environmental Biology, Utrecht University, Utrecht, the Netherlands (Joy Debnath).

## Abstract

Salt stress alters plant development, including the floral transition, but regulation of timing of flowering by salt is poorly understood at the molecular level. To identify genetic loci regulating the floral transition under high soil salinity, we performed a genome-wide association study (GWAS) in *Arabidopsis thaliana* and identified natural variation at the *UGT74E1-UGT74E2-BT3* (*UUB*) locus that correlates with bolting time specifically in response to salt stress. Genetic analysis revealed *BT3* as a novel repressor of the floral transition in control conditions. Similarly, the putative IBA glycosylases *UGT74E1 & UGT74E2* delay the floral transition in control conditions. Furthermore, we identified that IBA homeostasis regulators TOB1 and ECH2/IBR10 play a key role in the floral transition, and that ECH2/IBR10 are required for the early flowering phenotype of the *ugt74e1/ugt74e2* double mutant, indicating that UGT74E1 & UGT74E2 delay flowering by altering IBA homeostasis. A pangenome analysis of the *UUB* locus revealed variation in the occurrence of the DNA transposon *SAUERKRAUT* (*SKRT*). CRISPR-mediated *SKRT* deletion in Col-0 affected gene expression both within and outside the *UUB* locus and caused a salt-dependent delayed floral transition. The delayed bolting phenotype of the *skrt-2* mutant also depends on *ECH2/IBR10* function, indicating that *SKRT* accelerates the floral transition by altering IBA homeostasis. Finally, targeted demethylation of *SKRT* resulted in delayed floral transition under salt stress. Taken together, our data show a role for *SKRT* and its DNA methylation levels in the salt-dependent bolting time response in Arabidopsis, revealing a novel molecular mechanism to control flowering in adverse conditions.

## Introduction

Soil salinisation is an emerging problem, causing widespread soil degradation, consequently reducing crop yield (Daliakopoulos *et al*., 2016; van Zelm *et al*., 2020). In plants, salt induces ion toxicity and osmotic stress, significantly hindering crop growth and altering plant development, including root system architecture and the floral transition (Cho *et al*., 2020; van Zelm *et al*. 2020). The floral transition marks the shift from the vegetative phase to the reproductive phase and is a highly adaptive transition in a plant’s life cycle (Kinoshita & Richter, 2020). This transition is characterized by differential cell division in the shoot apical meristem (SAM), during which the SAM switches from producing leaves to producing inflorescences and flowers. This is followed by elongation, or bolting, of the flower-bearing inflorescence (Kinoshita *et al*., 2020). Timing of flowering enables plants to synchronize internal reproduction cues, *e*.*g*. age & gibberellic acid (GA) levels, with favourable environmental cues, such as photoperiod & temperature. This balancing act enhances the likelihood of successful fertilisation and seed development and is thus key for plant fitness and ensures high agricultural yields (Purugganan and Fuller, 2009; Kinoshita & Richter, 2020).

Although it remains unclear how salt is perceived by plants, salt has been found to influence several floral transition pathways in Arabidopsis, including the photoperiod, GA and age pathways (van Zelm *et al*. 2020; Kazan & Lyons, 2015; Jiao *et al*., 2024). In the photoperiod pathway, in long days (LD) the circadian clock regulates flowering mainly by targeting the protein GIGANTEA (GI) that stimulates the accumulation of CONSTANS (CO) protein. CO in turn activates the expression of the florigen *FLOWERING LOCUS T (FT)* in phloem companion cells (Kinoshita & Richter, 2020). Subsequently, FT travels to the shoot apical meristem (SAM) and binds the transcription factor FLOWERING LOCUS D (FD; (Gao *et al*., 2025). The FT-FD module then induces the expression of floral meristem identity genes and shifts the SAM to form a reproductive meristem that gives rise to inflorescence- and floral meristems (Kinoshita *et al*., 2020; Kinoshita & Richter, 2020). High salt thwarts this process, inducing the degradation of GI by the 26S proteasome, leading to reduced *CO* and *FT* expression and the production of BROTHER OF FT AND TFL1 (BFT) that competes with FT for the interaction with FD (Li *et al*., 2007; Kim *et al*., 2013; Ryu *et al*., 2011; Ryu *et al*., 2014). This ultimately leads to a delay of the floral transition in salt.

To identify genetic loci regulating the floral transition upon salt stress beyond known flowering regulation pathways, we performed a genome-wide association study (GWAS) in *Arabidopsis thaliana*. This GWAS revealed several loci that correlate with timing of floral transition in response to salt stress, and we focussed on the *UUB* locus that contributes to regulation of salt-dependent bolting time. Genetic analysis revealed that presence of the *BT3* and the putative indole-3-butyric acid (IBA) glycosylases *UGT74E1 & UGT74E2* genes at this locus delay the floral transition in control conditions. We found the *SAUERKRAUT (SKRT)* DNA transposon in the *UGT74E2-BT3* intergenic region to be responsible for accelerating bolting and flowering time in salt conditions. In line with this, we find clock (- regulated) genes and *BT3* differentially expressed in *SKRT* deletion mutants. Moreover, CRISPR-mediated DNA demethylation of *SKRT* indicate that its DNA methylation status plays an important role in bolting and flowering time regulation under saline conditions.

## Results

### Natural variation in salt-dependent floral transition phenotypes of Arabidopsis accessions

To investigate natural variation in the floral transition in response to salt stress, we recorded bolting time, flowering time and rosette leaf number (RLN) in LD conditions in *Arabidopsis thaliana* (*At;* Arabidopsis) accessions from the HapMap population (Table S1; Weigel & Mott, 2009). These 97 accessions – selected against vernalization requirement - were tested in control and salt stress conditions in soil, where salt treated plants were given a single watering of NaCl at 7 days after sowing (DAS). Within this population we identified variation in bolting and flowering time and RLN in control & salt conditions and in responsiveness to salt (Figure 1A, S1A&B, Table S1).

**Figure 1.**
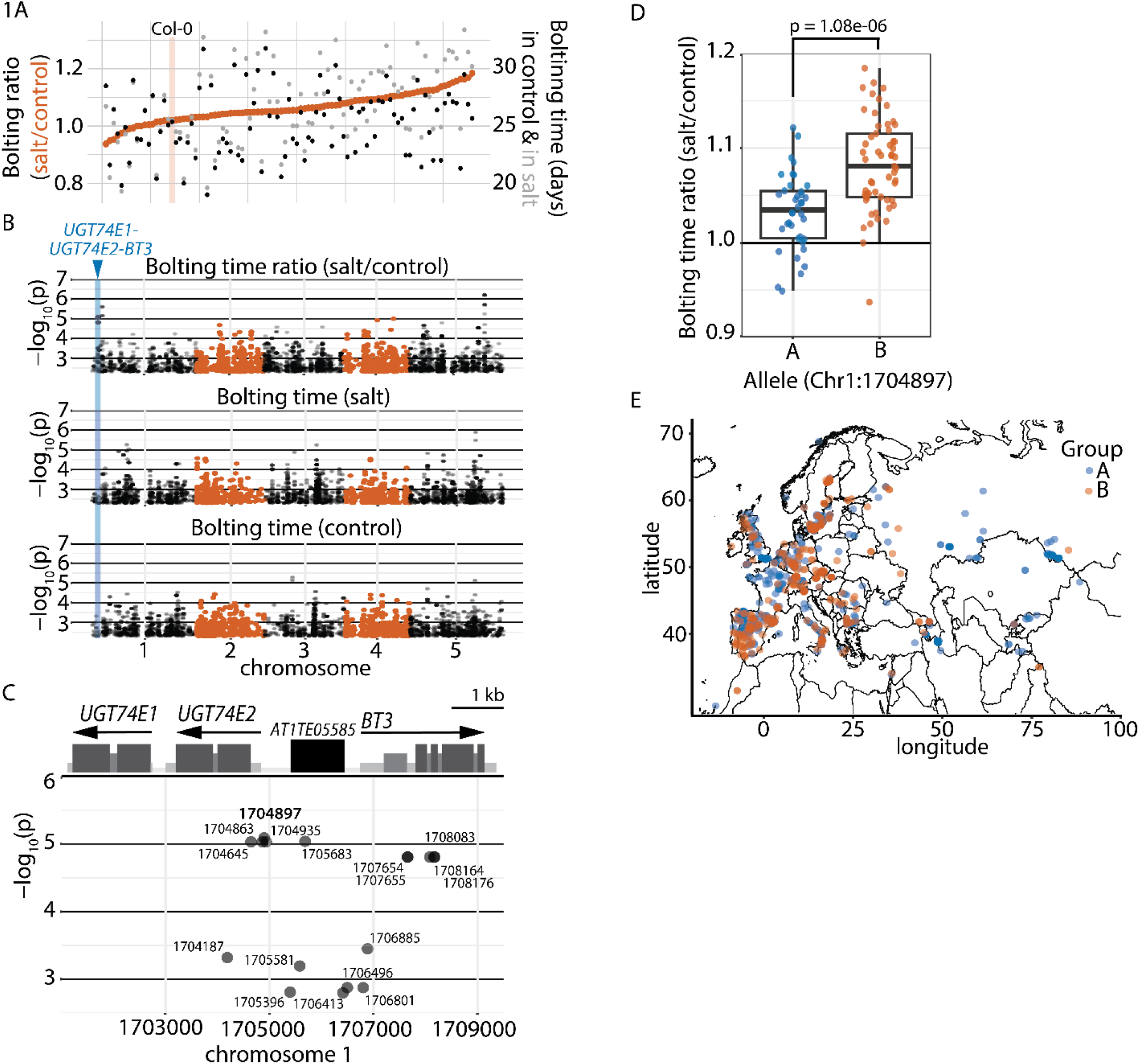
*Arabidopsis thaliana* natural variation in bolting time in response to salt correlates with variation at the *UUB* locus A. Natural variation in bolting time in control (black), salt (grey) and in bolting ratio (salt/control; orange) for tested accessions listed in Table S1 in climate-controlled greenhouse conditions with supplemented light. B. Manhattan plots for bolting time ratio (salt/control), salt and control conditions (See Table S2), *UGT74E1-UGT74E2-BT3* locus is highlighted in blue. C. Genomic position of significantly associated SNPs at the *UUB* locus in the tested population (See Table S2). D. Bolting time ratio (salt/control) for accessions used in this study carrying allele A (Col-0 variant) or allele B (non-Col-0 variant) at Chr1:1704897. E. Geographic distribution of most significant SNP (Chr1:1704897) at *UUB* locus represented by allele A (Col-0 type) and allele B (Non Col-0 type).

To identify candidate loci explaining natural variation in the floral transition in response to salt stress, we performed a GWAS using the mean of each trait for control, salt and the relative response to salt, *i*.*e*. ratio: salt/control (Figure 1B, S1C&D, Table S1). We focused on the relative salt response for which we observed several loci with significant SNPs (p<1e-05; allele frequency (AF) >20%; Figure 1B & Table S2). Within these quantitative trait loci (QTLs) we identified promising candidate genes (Figure 1B, S1, Table S2), among others, *WITH NO LYSINE KINASE1* (*WNK1*; AT3G04910; Saddhe *et al*., 2021) for relative flowering time, and *UDP-GLYCOSYL TRANSFERASE 74E1 & E2* (*UGT74E1 & UGT74E2*; AT1G05675-80) & *BTB AND TAZ DOMAIN PROTEIN* 3 (*BT3;* AT1G05690) for relative bolting time.

We chose to study the *UGT74E1-UGT74E2-BT3* (*UUB)* locus (bolting time ratio) in more detail, since *UGT74E1 & UGT74E2* gene expression is induced by salt (Tognetti *et al*., 2010). Moreover, ectopic overexpression of *AtUGT74E2* has been shown to delay bolting time in Arabidopsis and induce salt tolerance in rice seedlings (Tognetti *et al*., 2010; Wang T. *et al*., 2020). In addition, Arabidopsis *bt3* mutant seeds germinate slower in salt and show reduced root growth in salt (Saputro *et al*., 2023).

The most significant SNPs for relative bolting time in salt at the *UUB* locus reside in the promoter and the 5’ UTR of *UGT74E2*, and in a DNA transposon (AT1TE05585 in TAIR10; Figure 1C, Table S2). When we divide the population into two groups based on this most significant SNP in *pUGT74E2*, we observe a significant delay in relative bolting time for group B with the non-reference variant (Figure 1D), suggesting this locus affects the bolting response upon salt stress. We did not observe a correlation between the SNP and the geographic distribution within the 1001 genomes collection (Figure 1E).

### *BT3* delays bolting and flowering time in control conditions

Since two significant SNPs reside inside the *BT3* coding region (K>N & N>K substitutions; Figure 1C), we first investigated the role of *BT3* in bolting time regulation. *BT3* encodes for one out of five BT proteins found in Arabidopsis, a family that is only found in plants and is defined by the fusion of a bric-a-brac, tramtrack & broad (BTB) domain, a transcriptional adapter zinc finger (TAZ) domain and a calmodulin binding domain (Du & Poovaiah, 2004). *BT3* acts redundantly with *BT2* and *BT1 & BT2* in, respectively, male and female gametophyte development. Furthermore, *bt2/bt3* double mutants are gametophytic lethal, highlighting a crucial role of *BT3* in plant development (Robert *et al*., 2009).

To assess the role of *BT3* in bolting time regulation, we generated *bt3* mutants in Col-0 using CRISPR-iCas9z (Figure 2A & S2A; Stuttmann *et al*., 2021). We tested these *bt3-crispr (bt3-c)* mutants for floral transition phenotypes in control and salt in LD conditions. Two independent frame shift mutants *bt3-c1* and *bt3-c2* bolted and flowered significantly earlier relative to Col-0 in control conditions, but not in salt (Figure 2B-C & S2B). No difference in RLN was observed between the genotypes (Figure S2C). To assess whether the difference in bolting and flowering time of these mutants was caused by a delay in the floral transition at the shoot apical meristem (SAM), we studied the morphology of the SAM at 14, 18 and 22 DAS in Col-0 and the *bt3* mutants, but did not observe marked differences between the genotypes (Figure S2D-E). Taken together, these results indicate that *BT3* negatively regulates bolting and flowering time in Col-0 in control conditions, but not in salt and that the difference in timing arises after the transition has occurred at the SAM.

**Figure 2.**
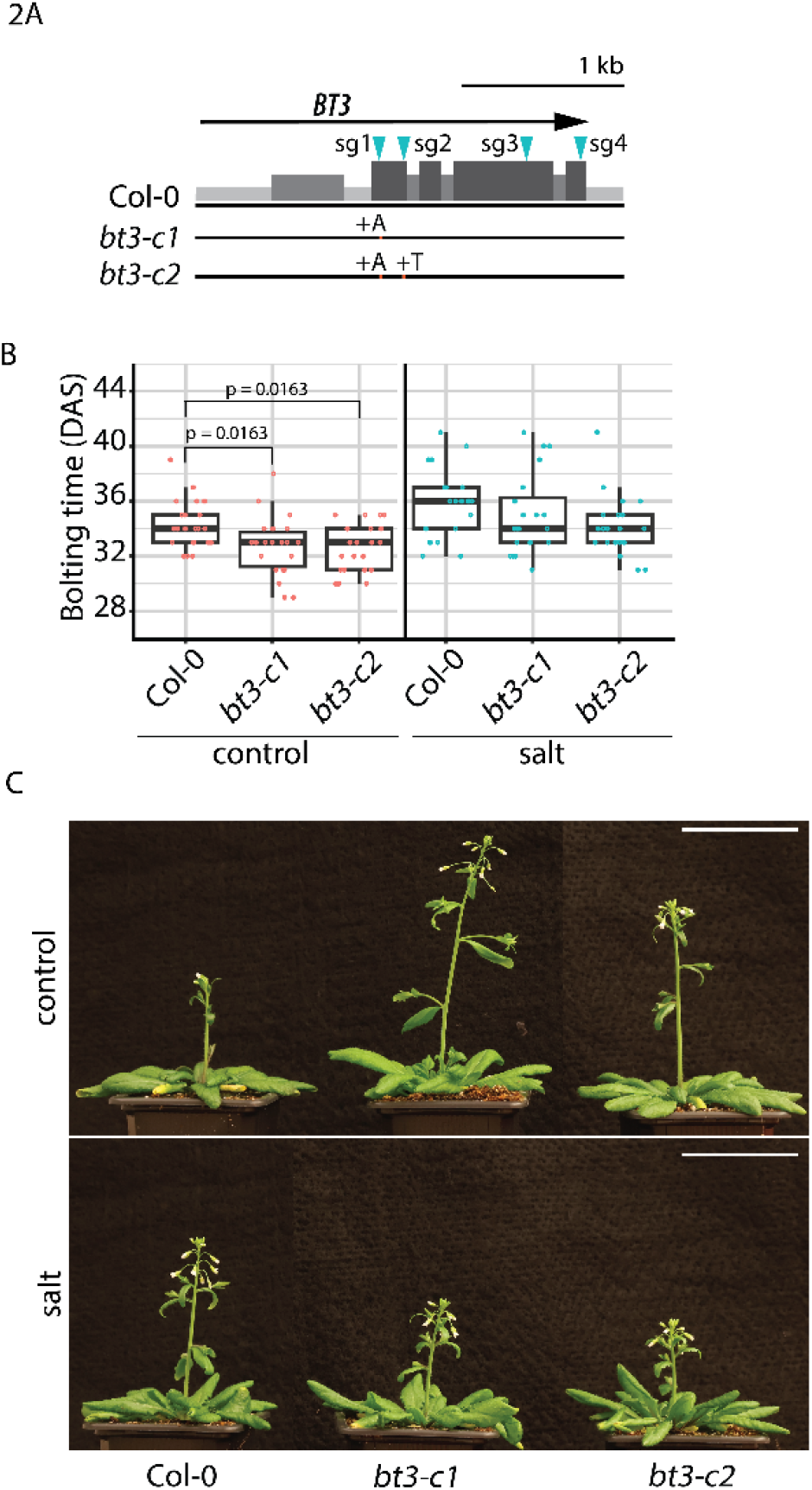
A. *BT3* delays bolting time in control conditions A. CRISPR mutagenesis strategy for *BT3* with sgRNAs (sg) binding sites in blue, UTRs in light grey, introns in grey and exons in dark grey. Letter and orange bar indicates DNA insertion and location (see Figure S2A for precise mutation). B. Bolting time for genotypes grown in climate-controlled growth chambers in LD in control (red) and salt (blue) conditions (Data are pooled from two independent experiments, n = 20 – 23). P-values within control or salt treatment indicate significant differences between genotypes using Pairwise Wilcoxon Rank Sum Test with Benjamini-Hochberg multiple testing correction. C. Representative image of shoot architecture at 39 DAS of genotypes shown in C, grown in LD in control and salt conditions. Scale bar = 5 cm.

### *UGT74E1 & UGT74E2* delay bolting and flowering time in control conditions

Next, we decided to investigate UGT74E2’s role in bolting time regulation, since previously UGT74E2 was shown to glycosylate the indole acetic acid (IAA) precursor indole butyric acid (IBA), and ectopic overexpression lines of *UGT74E2* delayed bolting and flowering (Tognetti *et al*., 2010). T-DNA mutants of *UGT74E2* did not show differential bolting or flowering time (Tognetti *et al*., 2010), possibly due to redundancy between UGT74E2 and its closest relative UGT74E1 (Figure 3A). This putative redundancy is supported by their 88% protein sequence similarity sequence (Table S3) and their highly similar protein structure, as modelled by Alphafold2 (Figure 3B, Video S1).

**Figure 3.**
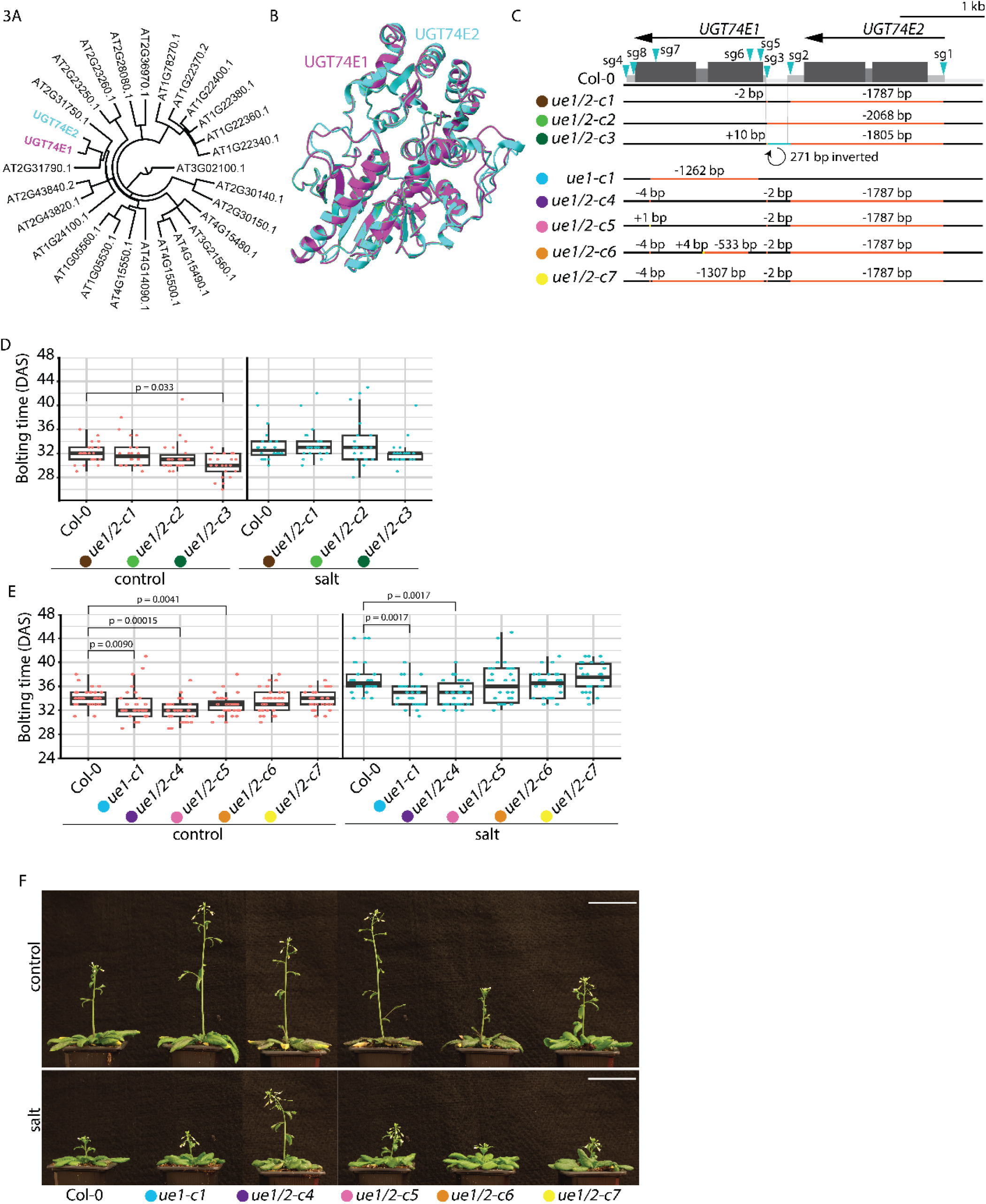
*UGT74E1* and *UGT74E2* delays bolting time in control conditions A. Phylogenetic tree of UGT74E2’s closest relatives in Arabidopsis Col-0. Relatives identified using BLAST-P aligned using MUSCLE and tree calculated using neighbour joining and a BLOSUM62 matrix. See Table S3. *B*. Overlap of AlphaFold protein structure models of UGT74E1 (magenta) and UGT74E2 (turquoise; Varadi *et al*., 2022). See Video S1 for a 3D rendering of the overlapped protein models. C. CRISPR mutagenesis strategy for *UGT74E1 & UGT74E2* with sgRNAs (sg) in blue, UTRs in light grey, introns in grey and exons in dark grey. Orange bar indicates DNA deletion; yellow bar indicates DNA insertion and blue bar indicates DNA inversion (see Figure S3A for precise mutation). Genotypes are colour coded, also in D-F. *ue1/2-c4-* to *-c7* were made in the *ue1/2-c1* background. D & E. Bolting time for genotypes grown in climate-controlled growth chambers in LD in control (red) and salt (blue) conditions (All data per plot is pooled from two independent experiments, D: n = 20 – 23; E: n = 26 – 31). P-values within control or salt treatment indicate significant differences between genotypes using Pairwise Wilcoxon Rank Sum Test with Benjamini-Hochberg multiple testing correction. See figure S3B-E for flowering time and RLN. F. Representative image of shoot architecture at 39 DAS of genotypes shown in F, grown in LD in control and salt conditions. Scale bar = 5 cm.

To investigate the role of *UGT74E1* and *UGT74E2* in bolting time regulation, we made *UGT74E1* and *UGT74E2 (UE1/2)* double mutants using CRISPR-iCas9z in Col-0 (Figure 3C; Stuttmann *et al*., 2021). We obtained one *ue1* mutant (*ue1-c1*) and seven *ue1/2* double mutants (*ue1/2-c1 to c7)*, all carrying a *UGT74E2* deletion and various *UGT74E1* promoter and *UGT74E1* promoter & CDS changes (Figure 3C & S3A). Three *ue1/2* double mutants, *i*.*e. ue1/2-c3/-c4/-c5*, bolted and flowered significantly earlier than Col-0 in control conditions, while in salt conditions only *ue1/2-c4* bolted and flowered earlier than Col-0 (Figure 3D&E & S3B&D). Furthermore, the *ue1-c1* single mutant bolted earlier in control and salt conditions and only flowered earlier in salt conditions (Figure 3E & S3D). None of the generated mutants showed an RLN phenotype (Figure S3C&E). These data suggest that *UGT7E1 & UGT74E2* delay the bolting time response in control conditions.

Given the differences in bolting time and flowering time between *ue1, ue1/2-c4 & -c5*, and *ue1/2-c6 & - c7*, we assessed whether the difference in bolting and flowering time of these mutants in control and salt conditions was caused by a clear delay in the floral transition at the shoot apical meristem (SAM). At 18 DAS in Col-0 only few SAMs had started to form inflorescence meristems, while *ue1-c1, ue1/2-c4, ue1/2-c6* and *ue1/2-c7* had already started to produce inflorescence meristems and sometimes even floral meristems (Figure S3F&G; Kinoshita *et al*., 2020, Cerise *et al*., 2023). This indicates that all *ue1/2* alleles accelerate the floral transition at the SAM, however, for *ue1/2-c6* and *ue1/2-c7* this does not manifest in an early bolting and flowering phenotype. Taken together, our observations indicate that *UGT74E1 & UGT74E2* delays the floral transition in control conditions. Furthermore, the phenotypes in salt, where only a selection of the mutants affect bolting and flowering in salt, suggests that there are complex interactions at this locus beyond the coding region of the UGT isoforms.

### Modulation of the floral transition by *UGT74E1 & UGT74E2* requires IBA

As *UGT74E1 & UGT74E2* regulate bolting and flowering time (Figure 3; Tognetti *et al*., 2010), and UGT74E2 functions in glycosylating the IAA precursor IBA, this suggests a regulatory role for IBA in the floral transition (Tognetti *et al*., Wang T. *et al*., 2020). To learn more about the role of IBA in the floral transition we studied the IBA homeostasis mutants *ech2-1/ibr10-1* and *tob1-1* (Michniewicz *et al*., 2019; Damodaran & Strader, 2024). The enzymes ENOYL-COA HYDRATASE2 (ECH2) and INDOLE3-BUTYRIC ACID RESPONSE1 (IBR10) convert IBA into IAA in the peroxisome (Strader *et al*., 2011; Damodaran & Strader, 2019). On the other hand, *TRANSPORTER OF IBA1 (TOB1)* is an IBA transporter in the vacuolar membrane that has been suggested to indirectly repress auxin availability, and thus auxin signalling (Michniewicz *et al*., 2019; Damodaran & Strader, 2019). We assessed the floral transition phenotypes for *ech2-1/ibr10-1* and *tob1-1* in control and salt conditions. In line with previously described root development phenotypes, *ech2-1/ibr10-1* and *tob1-1* showed contrasting phenotypes (Michniewicz *et al*., 2019). *Ech2-1/ibr10-1* bolted and flowered significantly later than Col-0 in control and salt, while *tob1-1* bolted significantly earlier than Col-0 in control and salt. (Figure 4A-C). Notably, only *tob1-1* showed a significantly lower RLN relative to Col-0 in control and salt (Figure S4A), which is reminiscent of auxin’s role in primordium outgrowth and phyllotaxis at the SAM (Galvan-Ampudia *et al*., 2020).

**Figure 4.**
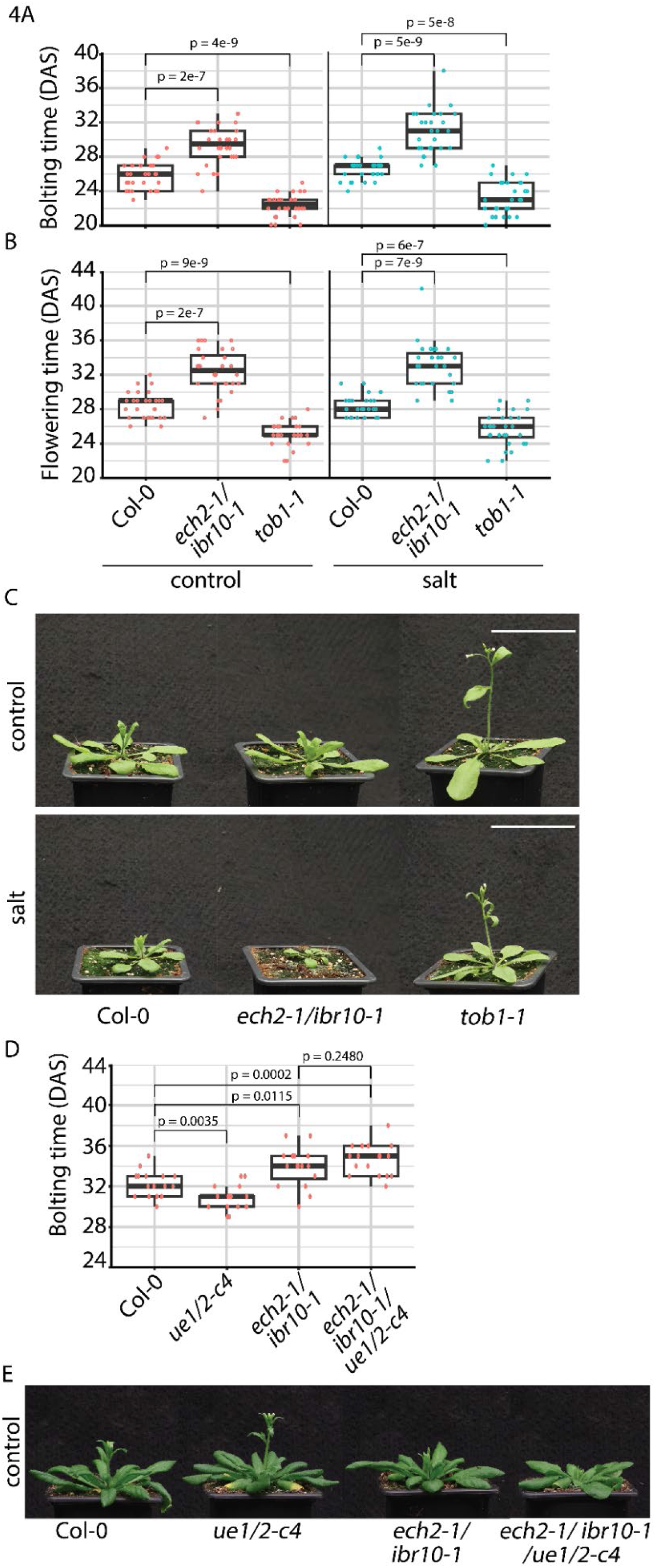
Bolting and flowering time regulation by *UGT74E1 & UGT74E2* requires IBA catabolic enzymes ECH2/IBR10 A & B. Bolting time and flowering time for genotypes grown in a climate-controlled greenhouse with supplemented light in LD in control (red) and salt (blue) conditions (Data are pooled from two independent experiments, n = 27 – 28). P-values within control or salt treatment indicate significant differences between genotypes using Pairwise Wilcoxon Rank Sum Test with Benjamini-Hochberg multiple testing correction. C. Representative image of shoot architecture at 25 DAS of genotypes shown in A & B, grown in a climate-controlled greenhouse with supplemented light in LD in control and salt conditions. D. Bolting time for genotypes grown in climate-controlled growth chambers in LD in control conditions (Data from one independent experiment, n = 16 – 17). P-values indicate statistical differences between genotypes using Pairwise Wilcoxon Rank Sum Test with Benjamini-Hochberg multiple testing correction. E. Representative image of shoot architecture at 31 DAS of genotypes shown in D, grown in LD in control and salt conditions.

To assess whether the *ue1/2-c4* early bolting and flowering mutant phenotype is dependent on IBA to IAA conversion, we crossed *ue1/2-c4* to *ech2-1/ibr10-1*. Next, we quantified the floral transition phenotypes for Col-0, *ue1/2-c4, ech2-1/ibr10-1* and *ue1/2-c4/ech2-1/ibr10-1* in control conditions. The *ue1/2-c4/ech2-1/ibr10-1* mutant phenocopied *ech2-1/ibr10-1*, by bolting and flowering significantly later than Col-0 (Figure 4D-E; S4B&C). This indicates that the *ue1/2-c4* accelerated floral transition phenotype is dependent on the IBA-to-IAA conversion by ECH2 and IBR10.

### The DNA transposon *SAUERKRAUT* regulates bolting time in Arabidopsis

As *UGT74E1, UGT74E2* and *BT3* all affect bolting and flowering time (Figure 2 & 3; Tognetti *et al*., 2010), we reassessed the natural variation at the *UUB* locus by retrieving the syntenic block from all published chromosome-scale genome assemblies of Arabidopsis accessions, as well as other species in the *Arabodae* supertribe (Table S4). We found that most structural variation between Arabidopsis accessions at this locus is due to presence or absence of an insertion of a REP11 family DNA transposable element (TE; AT1TE05585; Figure 5A). We named this REP11 TE *SAUERKRAUT (SKRT)* after the salty and sour/acidic fermented cabbage (Brassicaceae family), since this REP11 TE was identified in a salt experiment, adjacent to an indole-butyric acid regulating gene, in a Brassicaceae species. Notably, *SKRT* was not found in closely related *Arabidopsis* species or more distant Brassicaceae family members, indicating that the *SKRT* TE has been inserted during *Arabidopsis thaliana* evolution (Figure 5A; Table S4; File S1). Moreover, we do not observe a phylogenetic or geographic cluster containing the *SKRT* insertion (Figure 5B; S5), suggesting *SKRT* has been lost several times during *Arabidopsis thaliana* evolution. We noticed that *SKRT* was absent in 10 accessions that showed the strongest delay in bolting upon salt stress (Figure 5A & S5B and Table S1), which made us wonder whether *SKRT* may modulate the delay in bolting time upon salt stress.

**Figure 5.**
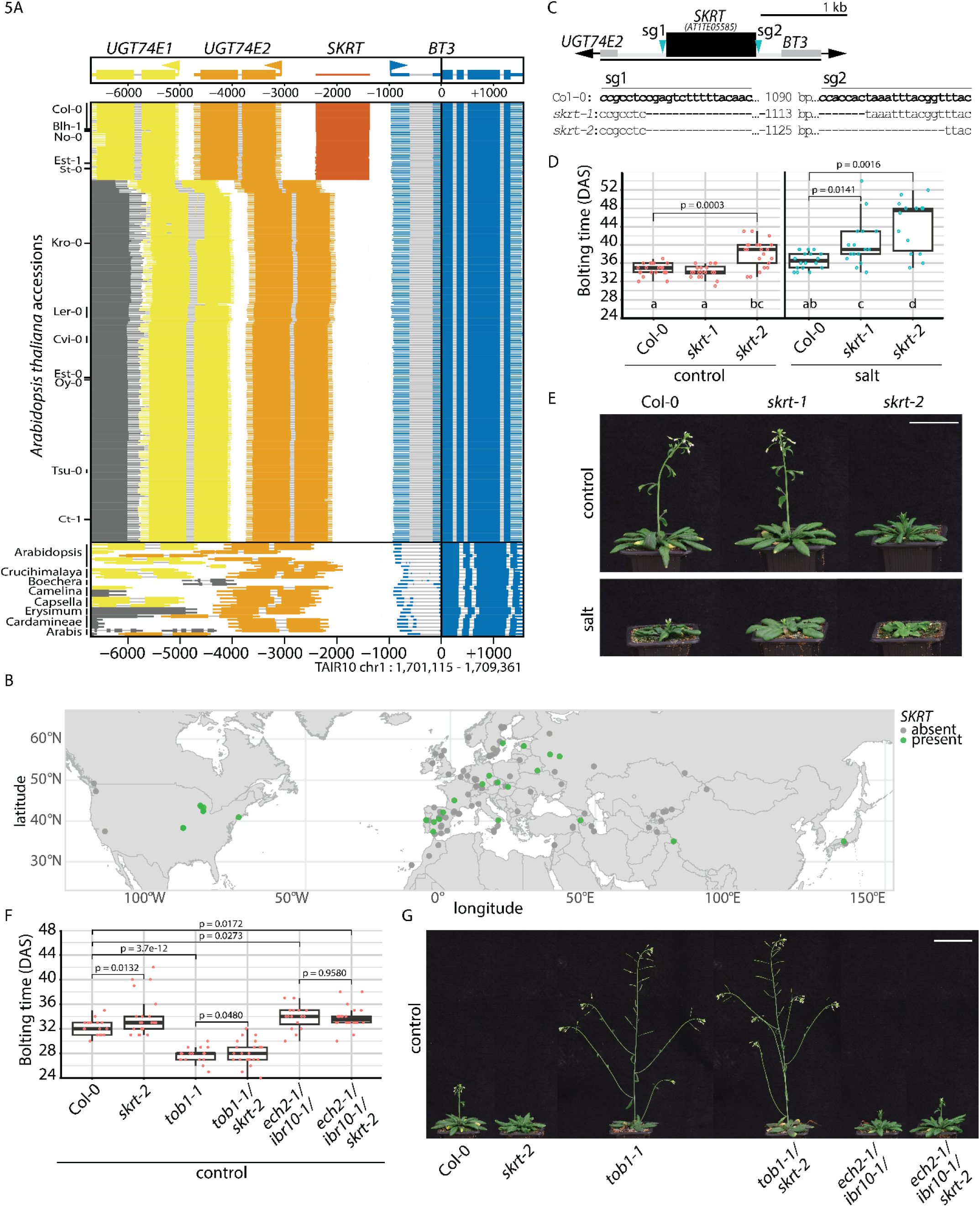
The DNA transposon *SAUERKRAUT (SKRT)* stimulates bolting time in control and salt. A. Alignment of the *UUB* locus in *Arabidopsis thaliana* accessions and closely related Brassicaceae species showing structural variation in the presence/absence of the *SKRT* DNA TE in the UGT74E2-BT3 intergenic region. Grey gene is UGT74E1’s upstream neighbouring gene. Indicated accessions were present in our natural variation screen, see Table S1. See File S1 for alignment of *UGT74E2-BT3* intergenic region. B. Geographic distribution of chromosome-scale Arabidopsis assemblies containing the *SKRT* locus. Green dots indicate accessions that contain the *SKRT* insertion, and grey dots are accessions where *SKRT* is absent. C. CRISPR mutagenesis strategy for *SKRT* with sgRNAs (sg) in blue, intergenic regions in light grey, UTRs in grey and *SKRT* in black. Alignment of *SKRT* target region in Col-0, *skrt-1* and *skrt-2* shown below diagram, where target site is shown in bold and PAM in italics. D. Bolting time for genotypes grown in climate-controlled growth chambers in LD in control (red) and salt (blue) conditions (Data from one independent experiments, n = 14-21, Figure S5E-G shows three more independent pooled experiments). P-values within control or salt treatment indicate significant differences between genotypes using Pairwise Wilcoxon Rank Sum Test with Benjamini-Hochberg multiple testing correction. Letters indicate significant (p < 0.05) differences between genotypes*treatments using a two-way ANOVA using Tukey-HSD multiple testing correction. E. Representative image of shoot architecture at 40 DAS of genotypes shown in D, grown in LD in control and salt conditions. F. Bolting time for genotypes grown in LD in control conditions (Data from two independent experiment, n = 33-43). P-values indicate statistical differences relative to control genotypes as determined by Pairwise Wilcoxon Rank Sum Test with Benjamini-Hochberg multiple testing correction. G. Representative image of shoot architecture at 40 DAS of genotypes shown in F, grown in LD in control conditions.

To assess whether *SKRT* could be responsible for the natural variation in bolting time upon salt stress, we generated *SKRT* deletion mutants in Col-0 using CRISPR-iCas9z (Stuttmann *et al*., 2021). These *SKRT* mutants, named *skrt-1 & skrt-2*, resemble natural *SKRT* deletions and lack the entire *SKRT* TE (Figure 5C). Next, we tested these lines for floral transition phenotypes in control and salt conditions. In contrast to Col-0, *skrt-1 & skrt-2* showed a significant salt-dependent delay in bolting and flowering time relative to *skrt-1 & skrt-2* in control (Figure 5C-D & S5C). This phenotype resembles accessions that delay bolting in response to salt. Furthermore, *skrt-2* showed a significant delay in bolting & flowering time in control and salt conditions compared to Col-0 (Figure 5D&E & S5C-G). We did not consistently observe significant differences in rosette leaf number for *SKRT* mutants in either control or saline conditions (Figure S5D&G).

Next, we assessed the morphological changes of the SAM during the floral transition in Col-0 and *skrt-2* at 14, 18, 22, 25 days after sowing (DAS) in control and salt conditions. We observed no differences between Col-0 and *skrt-2* in the timing of doming, production of reproductive meristems and production of floral meristems (Figure S5J&K). This suggests that *SKRT* does not play a major role in the induction of the floral transition at the SAM, but rather, *SKRT* regulates later stages of floral transition, *i*.*e*. outgrowth of the inflorescence and flower opening.

To assess whether the *skrt-2* phenotype requires IBA-to-IAA conversion for the delay in bolting time, we crossed *skrt-2* with *ech2-1/ibr10-1* and with *tob1-1* and grew these lines in control conditions. In line with previous experiments, *skrt-2* and *ech2-1/ibr10-1* bolted significantly later than Col-0, while *tob1-1* bolted significantly earlier than Col-0 (Figure 5F&G). Moreover, we observed that the *ech2-1/ibr10-1* can repress the *skrt-2* bolting time phenotype, while *tob1-1* cannot (Figure 5F&G & S5H&I). This indicates that *skrt-2* requires functional ECH2/IBR10 - and thus IBA-to-IAA conversion, to manifest a delay in bolting time.

### Clock-regulated & stress responsive genes are mis regulated in *skrt-2* shoot apices

Given that SAMs of *skrt-2* did not differ from Col-0 in control or salt conditions between 14-25 DAS (Figure S5J&K), we hypothesized that the bolting and flowering time phenotype is determined at the SAM between 22 DAS and 29 DAS, *i*.*e*. four days before the median bolting time of Col-0 in control (Figure S6A&B). To identify which of the genes in the *UUB* locus is regulated by *SKRT* and to find differentially expressed genes (DEGs) that can explain the bolting and flowering time delay in *skrt-2*, we employed RNA sequencing. For this, we extracted RNA from hand-dissected shoot apices from Col-0 and *skrt-2* at 22 DAS and 29 DAS grown in control and salt conditions (Figure S6A-C).Using DEseq2, we identified DEGs for genotype*treatment at 22 DAS and 29 DAS separately (Table S5&6). To determine differences during development we also assessed DEGs between 22 DAS and 29 DAS (Table S7-10). We further focused on the genotype*treatment contrasts, where at 22 DAS and 29 DAS we found between 61 and 526 *skrt-2-*regulated DEGs in control and salt conditions, and 5327 and 3697 salt-regulated DEGs in Col-0, respectively (p-adjusted <0.05; |LFC| > 0, Figure 6A&B; Table S5&6). We looked for intersections between these sets of DEGs (upset plots Figure 6A&B; Table S11&12) and for each intersection we looked for significantly enriched Gene Ontology (GO) terms (Table S13&14). We focussed on GO terms related to biological processes (BP) and found, among others, “rhythmic processes” and “response to stress” as the most significant GO terms for the overlaps between salt-regulated (Col-0) and *skrt-2-* regulated DEGs in control or salt (Figure 6A&B). Next, we looked at the expression patterns of genes within these GO terms and found several clock (-target) genes upregulated in Col-0 upon salt stress and in *skrt-2* in control conditions at 22 DAS and 29 DAS, such as *FKF1, PRR1, PRR3* and *PRR5*. In contrast, the clock-gene *LHY1* is downregulated in Col-0 upon salt stress and in *skrt-2* in control conditions at both time points (Figure 6C&S6D; Kinoshita & Richter, 2020). Furthermore, for the GO term “response to stress” at 29 DAS we find, among others, the floral repressor *JMJ30* upregulated by salt in Col-0 and is also *skrt-2* dependently upregulated in salt (Figure 6D; Gan *et al*., 2014). Next, we looked at the expression of *UGT74E1, UGT74E2* and *BT3* (Figure 6E).

**Figure 6.**
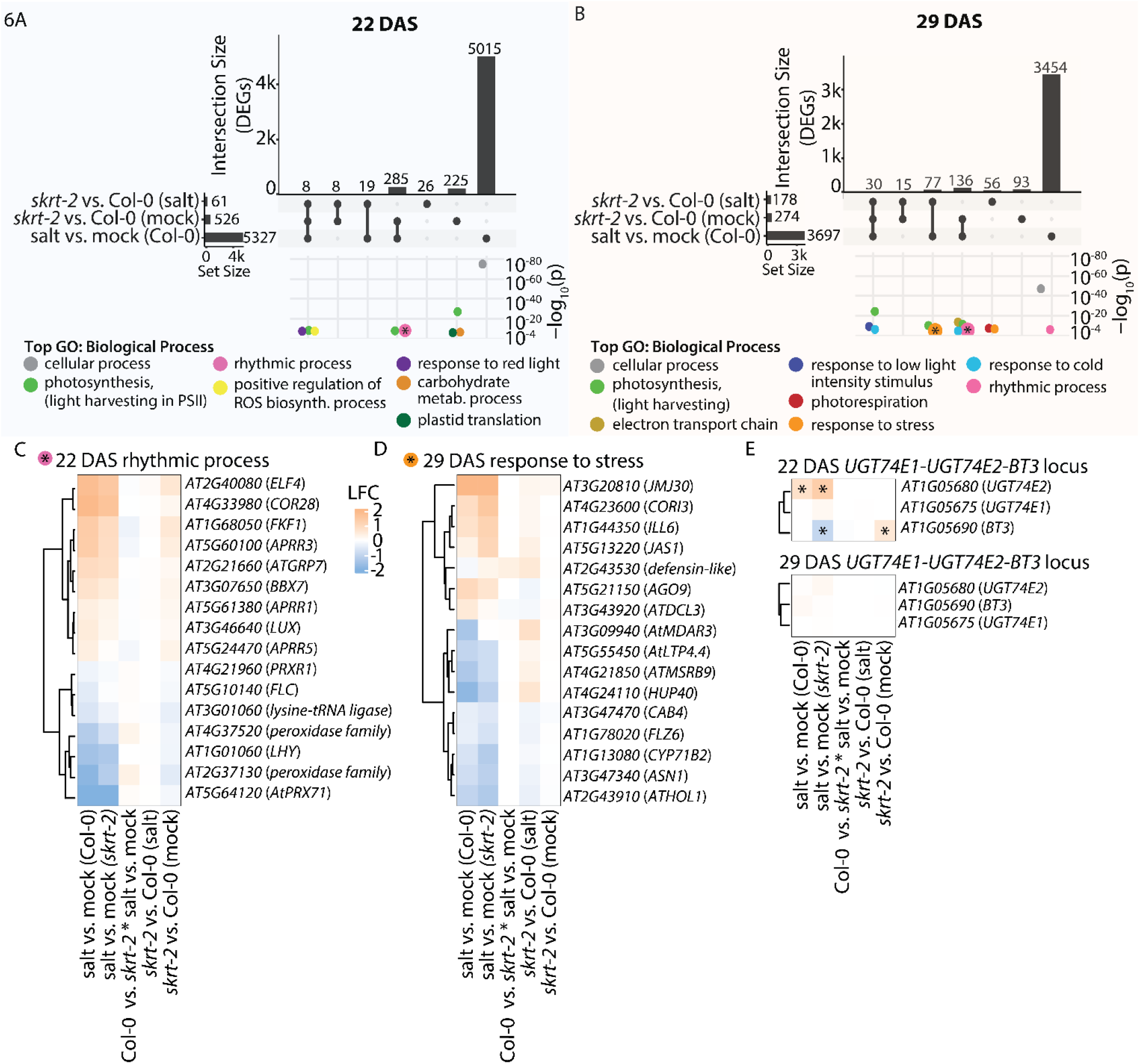
Salt-regulated genes in Col-0 and *skrt-2* regulated-genes function in rhythmic processes and response to stress. A & B. A: 22 DAS, B: 29 DAS. Top: Upset plot showing total number of DEGs per contrast and total number of overlapping genes between contrasts. Bottom: GO:BP enrichment analysis with on the x-axis the overlapping contrasts in upset plot and on the y-axis the -log_10_ p-value. P-value cut-off set to < -log_10_^-4^. GO:BP terms are colour coded. To reduce the number of redundant GO-terms we are only showing gProfiler’s “Highlighted” GO:BP terms (Kolberg *et al*., 2023), see S13&14 for the full list of GO terms. Within the GO term cellular process, several abiotic stress GO terms are represented, see Table S13 & 14. C-E. Heatmaps showing the relative genes expression as log_2_ fold change (LFC) for five contrasts of genes listed in C: 22 DAS GO:BP “rhythmic process” (See pink dot with asterisk in 6A), and D: 29 DAS GO:BP “response to stress” (See orange dot with asterisk in 6B) - with an LFC cut-off to reduce the number of genes (absolute LFC > 0.5, in at least one contrast), and E: the *UUB* locus at 22 and 29 DAS; asterisk indicates statistically significant difference for specified contrast (p < 0.05; DEseq2). See Table S5&6 for LFC values and p-adjusted values for all DEGs in all contrasts.

We observed significant upregulation of *UGT74E2* upon salt stress in Col-0 and *skrt-2* at 22 DAS. On the other hand, *BT3* was significantly upregulated in *skrt-2* relative to Col-0 in control conditions, but down regulated in *skrt-2* upon salt stress (Figure 6E). No significant differences in *UGT74E1* expression were found. In summary, these data indicate that salt affects the expression of, among others, clock genes in apices of Col-0. Moreover, these clock genes and *BT3* are also differentially expressed in *skrt-2* relative to Col-0.

### Demethylation of *SKRT* leads to a delay in bolting time upon salt stress

Since TEs can alter gene expression by introducing or disrupting *cis-*elements, but also by introducing RNA-directed DNA methylation (RdDM) to a locus (Hirsch & Springer, 2017; Tossolini *et al*., 2025), we investigated whether the DNA methylation pattern of *SKRT* would affect bolting time. We therefore assessed the DNA methylation at *SKRT* in Col-0 seedlings using a publicly available dataset and found that Col-0 *SKRT*, but not the coding regions surrounding *SKRT*, carries DNA methylation (Figure 7A; Fang *et al*., 2022). This means that *SKRT* mutants are not only *SKRT* DNA/*cis*-element deletion mutants, but also *SKRT* DNA methylation deletion mutants. Consequently, deletion of *SKRT* could therefore also alter the epigenetic landscape at the *UUB* locus. To assess whether *SKRT* methylation levels contribute to the *SKRT* mutant floral transition phenotypes, we set out to demethylate *SKRT*. To achieve this, we used a construct that expresses a catalytically dead-Cas9, the SunTag system and a C-terminal domain of the human demethylase TET1 (dCas9-SunTag-TET1; Figure 7B) as described by Gallego-Bartolomé *et al*. (2018).

**Figure 7A.**
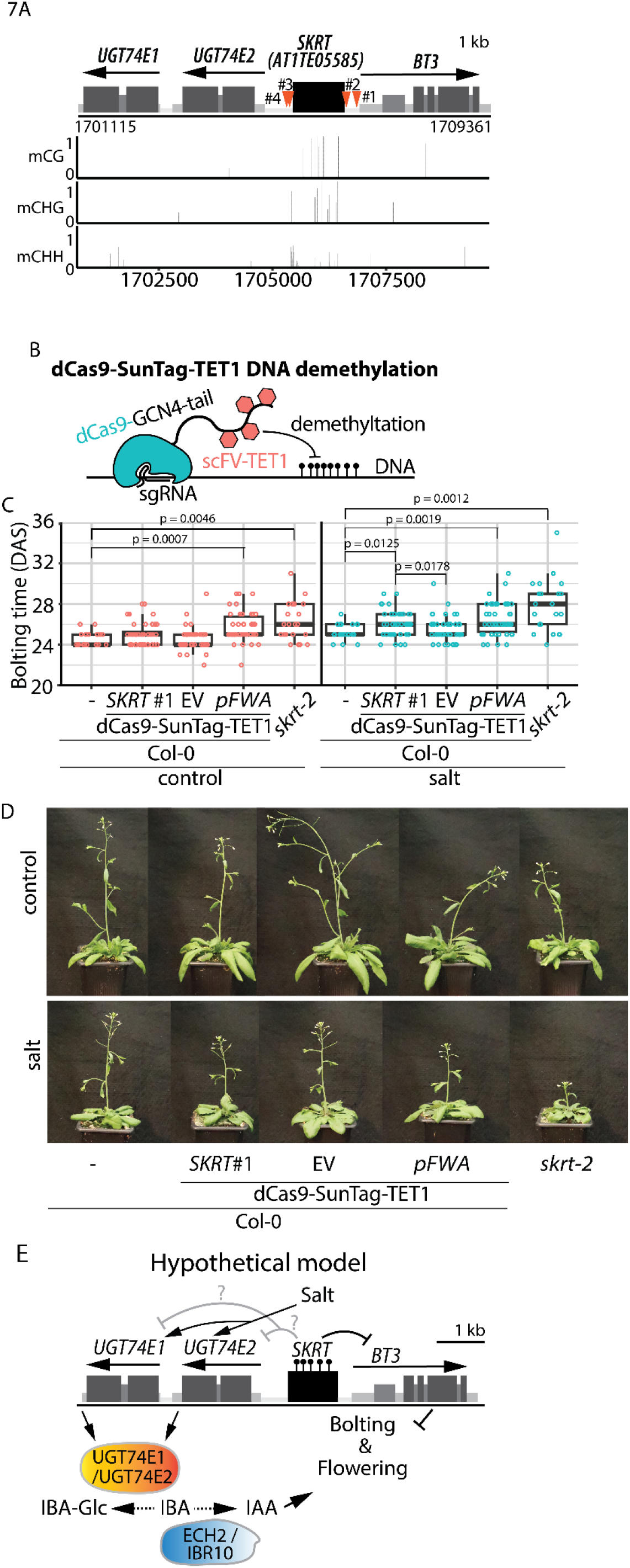
Demethylation of *SKRT* leads to a salt-dependent delay in bolting time. A. DNA methylation levels at the *UUB* locus for Col-0 in whole seedlings at 10 DAS (Fang *et al*. 2022). Red arrow heads and black numbers indicate the four sgRNA target regions expressed in the *dCas9-SunTag-TET1 SKRT* lines. B. Simplified schematic representation of the *dCas9-SunTag-TET1* DNA demethylation system (Gallego-Bartolomé *et al*. 2018). The sgRNA binds the target region and the dCas9 protein that is fused to an epitope tail consisting of ten GCN4 peptides. The scFv peptide that binds the GCN4 peptide is fused to sGFP and TET1 demethylase and several of these protein fusions (red) can bind the GCN4 tail. This GCN4-scFv dimer is called SunTag. This leads to several TET1-scFv protein fusions associating with the target region and removing DNA methylation (ball and stick icons) from the surrounding DNA. C. Bolting time for genotypes grown in a climate-controlled greenhouse with supplemented light in LD in control (red) and salt (blue) conditions. Data for *dCas9-SunTag-TET1 SKRT* lines #2-4 can be found in figure S7A-C. For Col-0 and *skrt-2* 21 replicates per genotype per treatment were used, for *dCas9-SunTag-TET1* lines 20 independent T2 families with 2-3 replicates per family were used per genotype per treatment (n = 44 - 59). P-values within control or salt treatment indicate significant differences between genotypes using Pairwise Wilcoxon Rank Sum Test with Benjamini-Hochberg multiple testing correction. D. Representative image of shoot architecture at 31 DAS of genotypes shown in C, grown in a climate-controlled greenhouse with supplemented light in LD in control and salt conditions. E. Working model of bolting time regulation at the *UUB* locus. *SKRT*, containing *cis*-elements and DNA methylation levels (represented by stick and ball symbol), stimulate bolting time and flowering time through the repression of *BT3* expression, but possibly also of *UGT74E1 & UGT74E2* (grey arrows). *BT3, UGT74E1 and UGT74E2* delay flowering and bolting. It remains unclear how *BT3* regulates the floral transition on a molecular level. UGT74E1 and UGT74E2 are putative IBA-glycosylases forming IBA-Glc from IBA, thereby diminishing the pool of free IBA that can be metabolized to IAA by ECH2/IBR10, which is required for timely flowering and bolting. Mutation of *ECH2/IBR10* is sufficient to repress the *ue1/2-c4 and skrt-2* floral transition phenotypes, indicating that *SKRT* requires IBA-to-IAA conversion to stimulate flowering and bolting time. Moreover, this suggests that *SKRT* represses *UGT74E1* and/or *UGT74E2* expression and thus their repressive effects on the floral transition (grey arrows). Based on the natural variation phenotypes and the *skrt-1, skrt-2* and *dCas9-SunTag-TET1 SKRT* line #1 phenotypes, we suspect that there is a salt-responsive regulation mechanism present in the *UGT74E2-BT3* intergenic region that can stimulate the expression of *UGT74E1, UGT742* and/or *BT3* in *SKRT* null accessions, leading to a delay in bolting time. Since we observed salt-dependent upregulation of *UGT74E2*, we suspect that this gene is differentially regulated by *SKRT* during the floral transition, however, we did not observe such *SKRT-*dependent *UGT74E2* regulation in the tissue or timepoints we analysed in our transcriptomic analysis. We hypothesize that in *SKRT*-containing accessions such as Col-0, *SKRT* can inhibit/diminish the salt-dependent upregulation of *UGT74E1, UGT742* and/or *BT3*, which leads to a limited or no delay in bolting time upon salt exposure.

We transformed Col-0 with several independent dCas9-Suntag-TET1 constructs: four constructs carrying different guides that target the upstream or downstream region of *SKRT* (Figure 7A) and as a negative control an empty vector carrying no guide. Furthermore, as a positive control we expressed a guide that targets the promoter of *FLOWERING WAGENINGEN* (*FWA)*, which was previously shown to delay the floral transition (Koornneef *et al*., 1991; Gallego-Bartolomé *et al*. 2018). To negate T-DNA insertion effects, we selected 16-20 independent T2 families per construct and scored their bolting & flowering time in control and salt conditions in LD.

Col-0 plants expressing *dCas9-SunTag-TET1 SKRT guide #1* showed a significant delay in bolting and flowering time in salt conditions relative to Col-0 and the empty vector control (EV) in salt conditions, but not in control (Figure 7C&D; S7A-C). This suggests that DNA methylation of *SKRT* is required to stimulate bolting upon salt stress. In line with previous experiments, the positive controls *skrt-2* and the *FWA-*targeting construct showed a significant delay in the timing of bolting and flowering in control and salt conditions (Figure 7C; S7A-C). In contrast, the empty vector control and the three remaining *SKRT* targeting constructs showed no significant delay in bolting time or flowering time compared to Col-0 in control or in salt (S7A-C). We verified these results by replicating the experiment with 4 independent *dCas9-SunTag-TET1 SKRT guide#1* T2 families for which we again observed a significant delay in bolting and flowering time compared to Col-0 in salt conditions, but not in control (Figure S6). Taken together, these data suggest that *SKRT* methylation is required to indirectly stimulate bolting time in Col-0 specifically upon exposure to saline conditions. Moreover, this CRISPR-demethylation approach independently confirmed that *SKRT* in the *UUB* locus regulates bolting time in a salt-dependent manner.

## Discussion

Soil salinisation is an emerging problem for global food safety, as salt significantly alters crop growth, development and the floral transition, thereby lowering the likelihood of successful fertilisation and seed development (Purugganan and Fuller, 2009; Daliakopoulos *et al*., 2016; Cho *et al*., 2020; van Zelm *et al*. 2020; Kinoshita & Richter, 2020). Several factors have been described to regulate the floral transition upon salt stress, which mostly seem to revolve around tweaking the photoperiod pathway components GI, CO and FT (Kazan & Lyons, 2015). Here we set out to identify novel loci that contribute to the floral transition upon salt stress.

Using a natural variation approach, we found several loci that play a role in the salt-dependent floral transition (Figure 1 & S1). We chose to investigate the *UUB* locus further and found that *BT3* mildly delays and flowering bolting time in control conditions, but not in salt (Figure 2 & 7E). Little is known about the molecular function of BT3 and its family members. In Arabidopsis, BT1 can interact with transcriptional regulators and bromodomain proteins BET9 and BET10 (Du & Poovaiah, 2004), and BT2 regulates telomerase activity (Ren *et al*., 2007). These functions align with their nucleocytoplasmic localisation. On the other hand, AtBT4 & AtBT5 localise solely to the cytoplasm (Robert *et al*., 2009). *AtBT3’s* closest relative, *OsBTBZ1* is also a nuclear localised protein and is upregulated upon salt stress in rice, moreover, *OSBTBZ1* lies in a salt tolerance QTL in rice (Saputro *et al*., 2023). These observations and our identification of *BT3* on a locus that affects salt-dependent bolting time in Arabidopsis, suggest that *BT3* could play a more general role in salt stress tolerance.

For mutants in *ugt74e1 & ugt74e2* we found an accelerated flowering and bolting time phenotype in control and salt conditions (Figure 3), where particularly mutations in *UE1 (ue1/2-4 & -c5)* or in a *pUE1* inversion (*ue1/2-c3*) on top of a full *UE2* deletion resulted in an accelerated floral transition (Figure 3). Given that *35S:UGT74E2* lines delay the floral transition (Tognetti *et al*., 2010), these observations together suggest that *UE1* is critical for the delay of the floral transition and that *UE2* acts redundantly with *UE1* (Figure 7E).

Since UGT74E1 & UGT74E2 are putative IBA glycosylases (Figure 3; Tognetti *et al*., 2010; Wang T. *et al*., 2020), we investigated the role of IBA in regulating the floral transition by studying IBA regulators TOB1 and ECH2 & IBR10. TOB1 and the combined action of ECH2 & IBR10, respectively, delay and accelerate the floral transition in control and salt conditions (Figure 4 & 7E). Moreover, we observed that the early bolting and flowering phenotype of the *ue1/2-c4* is repressed in the *ech2-1/ibr10-1* double mutant background (Figure 4), indicating that for *ue1/2-c4* to accelerate the floral transition ECH2/IBR10 need to convert IBA into IAA. This suggests that UGT74E1 & UGT74E2 glycosylate IBA to repress the floral transition (Figure 7E). This hypothesis is further supported by work from Tognetti *et al*. (2010) and Wang T. *et al*. (2020) who showed that AtUGT74E2 shows IBA glycosylase activity *in vitro* and *in planta*. The Erysimum, Cardamineae and Arabis clades lack *UGT74E1*, suggesting that *UGT74E2* was duplicated in the *Arabodae* supertribe, forming UGT74E1, giving rise to a protein that is highly similar to UGT74E2 (Figure 3A-B&5A; Table S3).

Given that *UGT74E1, UGT74E2* and *BT3* delay bolting and flowering time, we next investigated the TE *SKRT*. Based on our data, *SKRT* was introduced at the *UUB* locus during Arabidopsis evolution, and since then has been lost many independent times from Arabidopsis accessions from various geographic regions (Figure 5A & S5A). Notably, North American Arabidopsis accessions mostly contain *SKRT* (Table S4), and since wild Arabidopsis is invasive in the Americas, this indicates that the invasive founders of these populations are mainly *SKRT* containing. TEs are well known to affect gene expression, where increasing proximity of a TE to a gene correlates with a decrease in gene expression, and achieved by introducing or disrupting *cis-*elements, but also by introducing RNA-directed DNA methylation to a locus (Tossolini *et al*., 2025; Wang *et al*., 2013). For example, a recent study elegantly showed that *ELONGATION FACTOR-TU RECEPTOR (EFR)* expression is repressed by an inverted-repeat transposon downstream of *EFR* through the formation of a repressive chromatin loop (Mencia *et al*., 2025). This repressive effect of TEs on gene expression has been empirically shown for the rice salt-tolerance regulator *HIGH-AFFINITY K TRANSPORTER1;5 (OsHKT1;5)*, where the CRISPR-mediated deletion of a TE in *pOsHKT1;5* led to reduced *OsHKT1;5* expression and consequently reduced salt-tolerance (Wang *et al*., 2020). Similarly, we found that *SKRT* deletion and demethylation in Col-0 led to a salt-dependent delay in bolting and flowering time, phenocopying the most-delayed accessions in our natural variation screen (Figure 1, 5, S5 & 7). Since *BT3* was upregulated in *skrt-2* in control conditions, this indicates that *SKRT* represses *BT3* expression (Figure 6E). Together these observations in rice and Arabidopsis provide empirical evidence that TEs alter gene expression in plants (Tossolini *et al*., 2025; Wang J. *et al*., 2020).

Since the *ech2-1/ibr10-1* double mutant repressed the *skrt-2* mutant phenotype, we hypothesise that *SKRT* stimulates the floral transition by repressing the gene expression of the floral repressors *UGT74E1 & UGT74E2* (Figure 5F&G & 7E). However, *UGT74E1 & UGT74E2* were not differentially expressed in *skrt-2* shoot apices in our transcriptome analysis (Figure 6). Notably, light regimes were shown to affect *SKRT* methylation levels in leaves, which correlates with enhanced *UGT74E1 & UGT74E2* expression, but reduced *BT3* expression (Emmerson *et al*., 2025). This supports our hypothesis that *SKRT* also regulates *UGT74E1* and *UGT74E2* expression, but that we did not capture the right time or tissue to observe *SKRT*-dependent *UGT74E1* or *UGT74E2* expression changes.

In a previous natural variation study on root architecture changes in response to salt in Arabidopsis, several significant SNPs were identified of which 13 SNPs and their associated genes resided within 5000 bp from a REP11 TE (Julkowska *et al*., 2017; Tale S15) One of these SNPs is in close vicinity to the salt tolerance regulator *HKT1* (AT4G10310) that could act to restrict Na^+^ transport to the floral tissues (Garcia-Daga *et al*., 2025). This observation raises the question whether REP11 TEs may be under selective pressure to remain in genes that function in salt stress responses. This idea is supported by, among others, the observations by Deneweth *et al*. (2022), who found specific TE superfamilies enriched in genes that are differentially regulated upon heat stress in Arabidopsis and upon light stress in tomato (Tossolini *et al*., 2025). Since TEs make up 20-80% of crop genomes, they are the proverbial elephants in the genome and it is therefore worth going beyond annotating TEs, and increase our efforts to understand when, where and in which situation a transposon affects gene expression (Hirsch & Springer, 2017; Tossolini *et al*., 2025). In this study, we aimed to identify regulators of the floral transition in salt; consequently, we discovered a complex genomic locus in which multiple genes and a transposon tweak the timing of the floral transition in salt conditions. This opens new possibilities to fine-tune gene expression and consequently flowering, and plant resilience in general.

## Materials and Methods

### Plant materials, growth conditions and phenotyping assays

Arabidopsis thaliana plants of ecotype Columbia-0 (Col-0) were used in all experiments. The *ech2-1/ibr10-1* (Strader *et al*., 2011) and *tob1-1* (Michniewicz *et al*., 2019) lines were kindly provided by Lucia Strader. The CRISPR lines generated in this study, *i*.*e*., *skrt-1, skrt-2, ue1-c1, ue1/2-c1, -c2, -c3, -c4, -c5, -c6 & -c7, bt3-c1* and *bt3-c2*, were made in the Col-0 background, where *ue1/2-c4* to *c7* were made in the *ue1/2-c1* mutant background. The number of days until bolting was scored when the primary inflorescence reached a height of 1 cm measured from the first leaf’s petiole up to the tip of the top floral bud, flowering was scored upon opening of the flower, *i*.*e*. the stage in which fertilisation could take place, and rosette leaf number was counted after plants had bolted. Plots and statistics were generated in Rstudio using ggplot2 with occasional assistance from Mistral AI’s Le Chat (Wickham, 2016).

For plate-grown plants, *e*.*g*. for propagation and transformant selection, seeds were wet sterilized by 30% commercial bleach and 0.2% (v/v) triton-X 100 for 10 minutes and washed 5 times with sterile milliQ. Seeds were sown on ½ Murashige and Skoog (MS) medium including vitamins (Duchefa) containing 0.1% 2-Morpholinoethanesulfonic acid monohydrate (MES) buffer (Duchefa) and 0.8% plant agar (Duchefa) and pH was adjusted to 5.8 with KOH, and if applicable, with hygromycin B (15 μg/mL). Seeds were stratified on plate at 4 °C in the dark for 2-4 days. Plants were grown at 22°C and long day photoperiod (16 hours light, 120 μM/m^2^/s).

For the natural variation screen, 120 non-vernalisation requiring accessions from the HapMap collection (Weigel and Mott, 2009) were grown, out of which 95 accessions bolted (at least 5 plants in both control and salt conditions) during the course of the phenotyping experiment, and were used for further analysis (see Nordborg codes and mean values in Table S1). After seeds were allowed to germinate for 7 days on soil that was not watered with salt, seedlings were transferred to pots containing salt or control. For the salt treatment, 5×8 pots (Verspeentrays, Deens format, Desch-plantpak, NL) containing soil (Zaaigrond nr 1, SIR 27010-15, JongKind BV, NL) were dried for 4 days and subsequently watered from below with 75 mM NaCl (salt) or water (control) until the pots were fully saturated. After the initial salt treatment, all trays were watered twice a week from below with tap water. To prevent effects of location and tray, plants were transferred following a randomized block design. A week after transfer, additional seedlings were removed, leaving one seedling in each pot. The accessions were grown in the greenhouse of the University of Amsterdam in September and October 2016 (13 h – 11 h natural light), where long days was established by additional artificial light (16h light 20 °C / 8h dark 18 °C; relative humidity 60-80%).

For other soil grown plants (except the GWAS), seeds were stratified in demi-water 4 °C in the dark for 2-4 days and sown directly on standard potting soil. Seeds were covered for 3-4 days with a transparent plastic cover to achieve a high and consistent humidity level and germination. Genotypes were separated from each other in a Latin square design. Seven days after sowing, each tray was watered with 800mL of control solution (tap water, no suppletion of NaCl) or 800 mL salt solution that contained 18.40 g NaCl. Each tray contained 21 7×7 cm pots that can take up 3.5 L of water, where the water volume varied between ∼ 3 L to ∼4.3 L, meaning that the NaCl concentration varied between ∼75 mM and ∼105 mM NaCl. Afterwards, all plants were well-watered with tap water without added NaCl, to limit salt concentrations becoming too high and excluding water-limiting stress to affect plant phenotypes. Small-scale experiments were performed in controlled climate chambers with LEDs (LuxaLight LED-strip Neutral White 4300K Protected (24 V, 140 LEDs/meter, 2835 SMD, IP64; PAR (400-700nm): 130-150 μM/m2/s (400-500 nm: ∼35 μM/m2/s; 500-600 nm: ∼ 55 μM/m2/s; 600-700 nm: ∼ 50 μM/m2/s) set at long day (16h light 20 °C / 8h dark 18°C; RH: 50-70%), where light intensity and temperature gradually in-/decreased between ZT0-1 and ZT15-16. Large scale experiments were performed in the climate-controlled greenhouse (Wageningen University; Serre red 6.7-6.8) where long day was established by additional artificial LED light (16h light 20 °C / 8h dark 18 °C; relative humidity 60-70%) and these conditions resulted in a shorter flowering time compared to the controlled climate chambers. The climate controlled greenhouse experiments were performed between September 2024-October 2024 (12.4 h – 10.4 h natural light; dCas9-SunTag-TET1 T2 experiment big screen), November 2024-December 2024 (8.5 h – 7.5 h natural light; Col-0, *tob1-1* and *ech2-1/ibr10-1* experiment #1), and February-April 2025 (10.5 h – 14 h natural light; second dCas9-SunTag-TET1 T2 experiment & Col-0, *tob1-1* and *ech2-1ibr10-1* experiment #2).

### Genotyping

Plants were genotyped using Phire polymerase (ThermoFisher) using supplier’s settings (1’ 98 °C, 30x (10’’ 98 °C; 10’’ annealing; 10’’/kb elongation)) and oligos (ID Technologies; Table S16). In case of genotyping *SKRT*, PCR was performed using standard conditions other than an elongation step of 68°C for 2:30 to extend TA-rich regions (Su *et al*., 1996). For genotyping, (digested) PCR products were run on an 1% agarose gel and/or extracted using the PCR/Gel clean-up kit (Macherey-Nagel) using manufacturer’s protocols and sent for sanger sequencing (Macrogen). See Table S16 for primers and PCR & digest protocols.

### GWAS

Variant information for the used set of accessions was extracted from the 1.2M SNP data set based on the 1,001G Project filtered with cross-validation accuracy of 0.95 (Arouisse *et al*., 2020) and filtered for variants with a minor allele frequency > 0.05. The association analyses were conducted with GEMMA version 0.98 (Zhou and Stephens., 2012) using a univariate linear mixed corrected for population structure with a centred relatedness matrix calculated with GEMMA. To assess the geographic distribution of SNP 1:1704897 across Europe, the variant was extracted for all 1,001 and RegMap accessions from the same 1.2M SNP data set used to extract variant information for GWA analysis. The geographical distribution of both alleles was then plotted for European accessions using R packages ‘rworldmap’ and ‘rworldxtra’ (South, 2011).

### Alignments

UGT74E2-related protein sequences were obtained by BLAST search from Arabidopsis.org/tools/blast; TAIR10. Protein alignments were made with EBI’s MSA-MUSCLE

(www.ebi.ac.uk/jdispatcher/msa/muscle; Edgar, 2004), visualisation and tree calculation (Neighbour joining/BLOSUM62) were performed using JalView (Waterhouse *et al*., 2009), and trees were visualised using FigTree v1.4.4 (https://github.com/rambaut/figtree/).

#### Pangenome

Syntenic blocks were extracted from published *Brassicaceae* genomes using a MinHash sketch search. Briefly, all available genomes in the NCBI Genomes database included in the taxa “Brassicaceae” with a completeness of “chromosome” or above were downloaded. BinDash2 (Zhao *et al*. 2024) was used to create a minimizer sketch of each 100 kilobase window of each genome with 50kb overlap. The 2 blocks containing *UGT74E2* in the TAIR10 genome (GCF_000001735.4) were used as queries against all target blocks for the command *bindash dist -- mthres=0*.*2*. Of 396 input genomes, all target regions that contained consecutive hits to both query blocks were extracted as a FASTA file. One region was identified in each of 248 *A. thaliana* genomes, and one or more regions were found in an additional 39 species (Supplemental Table S4, File S1).

To annotate all syntenic blocks, Helixer was run on the resulting FASTA file via the Helixer web server (Holst *et al*. 2025). Protein sequences were extracted from all predicted genes and sequence similarity was determined in an all-vs-all configuration with DIAMOND v0.9.24 (Buchfink *et al*. 2021). To determine protein-coding gene orthologs, the all-vs-all DIAMOND results were clustered using the connected components algorithm of the igraph library in R (Csárdi & Nepusz 2006). To identify *SKRT* loci in the target blocks, BLASTN was used with the Col-0 *SKRT* sequence as a query.

Whole-genome phylogeny of the 287 genomes above was estimated by calculating all-vs-all genomic distance with Bindash2. This distance matrix was used as input to the FastME balanced Minimum Evolution Algorithm (Lefort *et al*. 2015). For spatial distribution of *SKRT* loci, latitude and longitude of all accessions were extracted from the AraPheno database (Togninalli *et al*. 2019) and plotted with the rnaturalearth package (Massicotte & South, 2026).

### Generating CRISPR mutants

Guide RNAs (gRNA) were selected using CHOPCHOP V3 (Labun *et al*., 2019), where target sites were only selected if off-targets contained >4 mismatches, which was double checked by BLAST (Phytozome: https://phytozome-next.jgi.doe.gov/; Araport11). Between 2-4 gRNAs per construct were cloned into *pAtU6-26* carrying shuttle vectors (pDGE332-337) and then to the transformation vector pDGE347 (*Basta Resistance*; *pOLE1::OLE1-RFP*; *pRS5a::intronised-ZmCas9::rbcS-E9 Terminator (Pisum sativum); 2-8x(pAtU6-26:gRNA)*) using the Stuttmann *et al*., 2021 cloning protocol. Plants were floral dipped (Clough & Bent, 1998) in Col-0, except *ue1/2-c4* to *-c7* that were dipped in the *ue1/2-c1* mutant background. Next, fluorescent T1 seeds were selected using a Leica MZ10-Fluorescent stereo microscope and grown on soil (Shimada *et al*., 2010). Mutations in T2 families were determined by performing DNA isolation on ∼40 T2 seeds for each T2 family (Cheung *et al*., 1993) and genotyping by PCR with primers spanning the CRISPR target region (See oligo list; Table S16). For T2 families that showed promising mutations, ∼8 non-fluorescent, *i*.*e*. non-transgenic, seeds were sown on soil and genotyped by PCR and sanger sequencing for mutations in the target region and the absence of the transgene was confirmed by PCR using the JS184 & JS185 primres or JAD022(M13f) primer and a reverse gRNA oligo that was used for cloning the vector (Table S16). If the transgene was still present in the selected mutant plant, the transgene was outcrossed or back-crossed to Col-0.

### Generating dCas9-SunTag-TET1

First, guides targeting *SKRT* surrounding region were cloned into the shuttle vector pDGE331 according to the pDGE cloning system protocol (Stuttmann *et al*., 2021). Next, we performed a PCR (Q5 polymerase; NEB) on this vector with primers JAD196+197 with a KpnI-HF overhang to amplify the pAtU6-26::sgRNA, followed by a KpnI-HF (NEB) digest and the PCR product was excised from gel using Macherey-Nagel PCR Clean-up, all according to standard manufacturer’s protocols. Next, we obtained the dCAS9-SunTag-TET1 vector, pEG302 and pEG302-FWA (Gallego-Bartolomé *et al*., 2018) from Gerco Angenent (WUR) and digested pEG302 with KpnI-HF (New England Biolabs (NEB), followed by a treatment with recombinant shrimp alkaline phosphatase (NEB) according to manufacturer’s protocols. This digested pEG302 vector was ligated with the pAtU6-26::sgRNA PCR product using T4 DNA ligase (NEB). The pEG302-*SKRT* vectors and pEG302-FWA were transformed in *Agrobacterium tumefaciens* AGL-0, dipped into Col-0 using floral dip and transformed seeds were selected on ½ MS media containing hygromycin B (15 μg/mL; Clough & Bent, 1998).

### RNA isolation and RNA-seq

Seven to nine apices per sample were harvested between ZT 7.5 and ZT 9.5 in triplicate and snap-frozen in liquid nitrogen and stored at -70 °C. Samples were homogenized using 2 stainless steel beads in 2 mL tubes (SafeSeal; Sarstedt) in a paint shaker (Fast & Fluid) until samples were fully homogenized. RNA isolation was performed using the Universal RNA purification kit (Roboklon; EURx, Berlin, Germany). Possible DNA contaminants were eliminated with an on-column DNaseI digest. Total RNA integrity and purity were assessed using an Agilent 5400 by Novogene (UK Co. Ltd (Cambridge, UK)) and subsequently used for RNA poly-A enrichment library preparation and transcriptome sequencing (150bp paired-end mode) on a NovaSeq 6000 platform (Novogene UK Co. Ltd (Cambridge, UK)). Reads were mapped to the TAIR10 genome (retrieved: 20240308) using the Nextflow (v. 23.04.3) rnaseq pipeline (rev. 3.9; Di Tommaso *et al*., 2017) yielding 15-75 million reads per sample. Next, genes with ≥2 transcripts per sample in all 3 replicates were filtered and significantly differentially expressed genes (p-adjusted (BH) < 0.05 ; |LFC| > 0) were assessed used DESeq2 (Love *et al*., 2014) and log fold changes were calculated using the “ashr” shrinkage estimation (Stephens, 2016). GO terms were assessed using gProfiler (Kolberg *et al*., 2023). Plots and statistics were generated in Rstudio with occasional assistance from Mistral AI’s Le Chat. Raw data has been reposited at EBI (https://www.ebi.ac.uk/biostudies/arrayexpress/studies/) and will be available online from April/May 2026.

### DNA methylation

DNA methylation data for 10-day-old Col-0 seedlings for the UGT74E2-BT3 region (chr1:1704640-1706807) were obtained from Fang *et al*., 2022 (GSE180635; https://ngdc.cncb.ac.cn/methbank/jbrowseGSE180635) and plotted in RStudio using ggplot2 with occasional assistance from Mistral AI’s Le Chat (Wickham, 2016).

### Protein structure

Protein structure models of UGT74E1 (Uniprot: P0C7P7) and UGT74E2 (Uniprot: Q9SYK9) were obtained from AlphaFold.com (Varadi *et al*., 2022) and visualised using PyMol v2.3.4.

### Confocal microscopy and imaging

Shoot apical meristem morphology was assessed at 14, 18, 22 and 25 DAS in soil-grown plants treated with control and 100 mM NaCl at 7 DAS, as described above for salt assays on soil (n = 3 - 7). Samples were hand-dissected to enrich for shoot apices by removing roots, leaves and leaf primordia and were then fixated for 20 minutes in PBS with 4% PFA under vacuum. Samples were then rinsed three times with PBS for 1 minute and incubated in ClearSee for whole tissue clearing for 1-3 weeks (Kurihara *et al*., 2015). Apices were stained for 1-4 hours with SCRI Renaissance stain (SR2200; 1:1000 in ClearSee, Musielak *et al*., 2015) to visualize cell walls and then imaged in water on a Zeiss LSM710 using a water dipping lens (Zeiss 40x/10 DIC VIS-IR), where SR2200 was excited at 405 nm and detected between 410-523 nm. SAMs were imaged as a Z-stack (thickness per slice = 3 µm) and were processed in ImageJ (Fiji; Schindelin *et al*., 2012). For image panels, each stack was processed to find the image representing the centre of the meristem. For analysis of inflorescence (i) and floral meristems (f), each stack was analysed separately. For the figures with whole plants, photos were taken using a Canon EOSR10 mirror-reflex camera with an RF 85 mm F2 macro lens or an RS-F 18-45 mm lens, where plants were kept at an equal distance from the lens for equal comparison between samples.

## Supporting information

Supl_File_S1_Fasta_Alignment

## Contributions

JAD, IK and CT conceived the project, JAD, AvD, RB and MAS generated plots and figure panels. IK, AJM and CT were responsible for the natural variation screen. RB, JAD and IK were responsible for the GWAS analyses. JAD and AMB performed PCRs on *SKRT*. JAD, PK, JD and SO cloned and selected the CRISPR mutants. AvD and JAD generated the crosses. JD generated the protein structure files. JAD and AvD performed SAM morphology experiments. MAS performed pangenome analysis. JAD and KD mapped and analysed the RNA-seq experiment. JAD and ETP cloned and selected dCAS9-Suntag-TET1 lines. JAD, TdZ, YHT and ETP performed flowering time assays. JAD and CT wrote the manuscript, with contributions and feedback from all authors.

## Data availability

RNA-seq data will be available on EBI (https://www.ebi.ac.uk/biostudies/arrayexpress/studies/) from April/May 2026 and code used to analyse and plot the data can be found on GitHub: https://github.com/DongusJA/SKRT_manuscript

## Acknowledgements

We would like to thank all staff in the greenhouses & labs for their support and all the BSc & MSc students at University of Amsterdam and Wageningen University & Research who contributed to this research as part of their course work or thesis. Furthermore, we would like to thank Lucia Strader for providing *tob1-1* and *ech2-1/ibr10-1*, Johannes Stuttmann for providing the pDGE vectors, Gerco Angenent for sharing pEG302 and pEG302-*pFWA* and Richard Immink for critical discussions.

## Competing interest

None to declare.

## Funding

This research was supported by funding from the Dutch Research Council (NWO), “Plant stress resilience: Think global, act local” (VI.C.192.033) to CT. MAS is supported by the NWO project “Lost in Translation” (VI.Veni.222.212).

## Supplementary data

Table S1. Natural variation in floral transition phenotypes: bolting time, flowering time and rosette leaf number (RLN) for tested HapMap accessions

Table S2. GWAS output for median values in Table S1

Table S3. Protein alignments for UGT74E2 & UGT74E1 and closest relatives of UGT74E2 in Arabidopsis

Table S4. Sequence data used to generate alignment of the *UUB* locus shown in Figure 5A

Table S5. DEG data for 22 DAS between for contrast: s*krt-2* vs. Col-0 and salt vs. control

Table S6. DEG data for 29 DAS between for contrast: s*krt-2* vs. Col-0 and salt vs. control

Table S7. DEGs in Col-0 in control conditions between 29 vs. 22 DAS

Table S8. DEGs in Col-0 in salt conditions between 29 vs. 22 DAS

Table S9. DEGs in *skrt-2* in control conditions between 29 vs. 22 DAS

Table S10. DEGs in *skrt-2* in salt conditions between 29 vs. 22 DAS

Table S11. Intersections between contrast for 22 DAS between s*krt-2* vs. Col-0 and salt vs. control

Table S12. Intersections between contrast for 29 DAS between s*krt-2* vs. Col-0 and salt vs. control

Table S13. GO enrichment for contrasts at 22 DAS Table S14. GO enrichment for contrasts at 29 DAS

Table S15. SNPs identified in Julkowska *et al*. (2017) for indicated phenotypes, where a REP11 lies within 5000 bp from the SNP

Table S16. Oligos used in the study for cloning, sequencing and PCR purposes

Video S1. Video showing an overlap between the UGT74E1 and UGT74E2 protein models.

File S1. Alignment of the *UGT74E2-BT3* intergenic region

## Supplemental Figures

**Figure S1.**
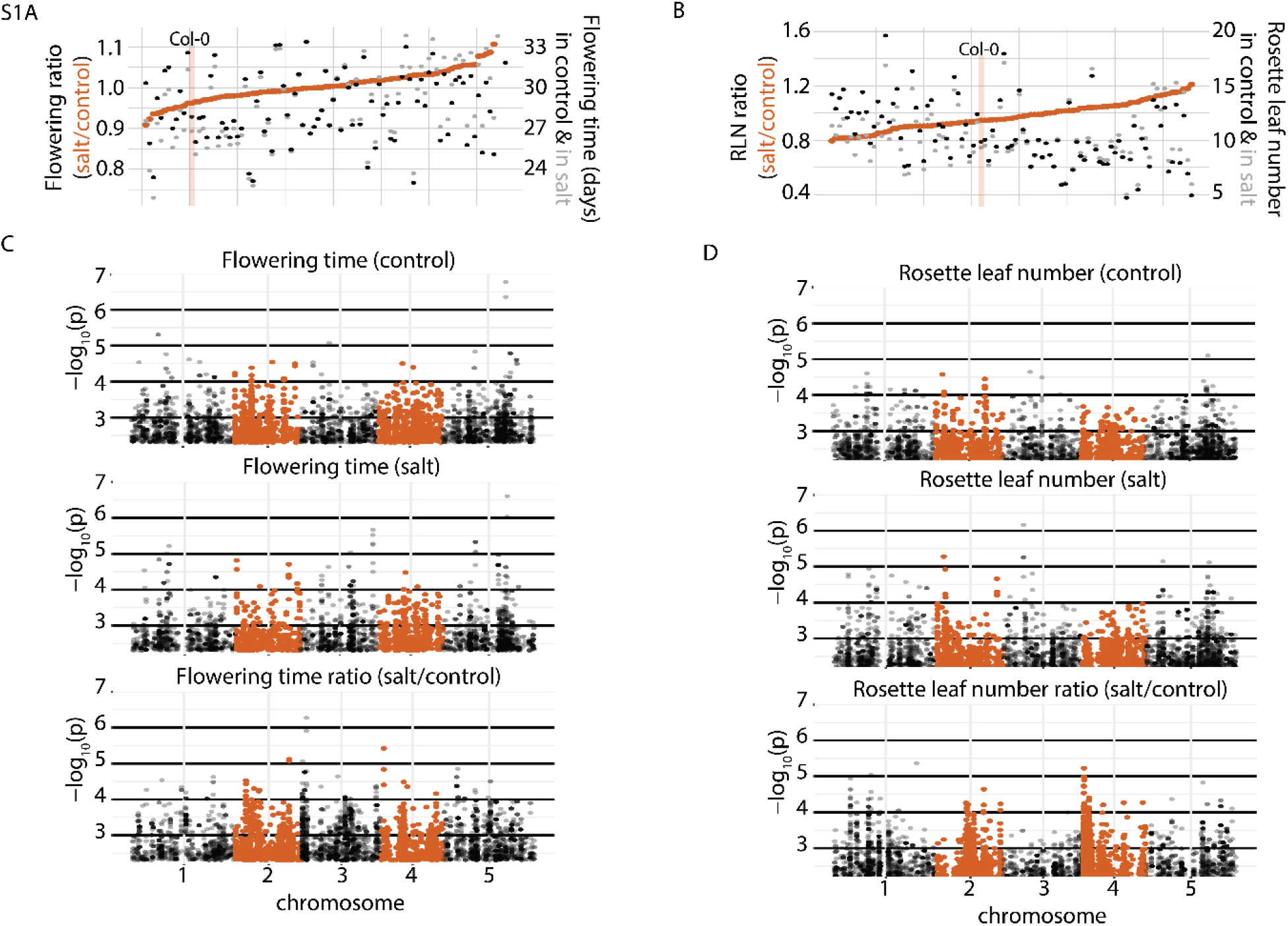
Natural variation in flowering time and RLN in control, salt and ratio (control/salt) A. Natural variation in flowering time in control (black), salt (grey) and in flowering time ratio (salt/control; orange) for tested accessions listed in Table S1. B. Natural variation in RLN in control (black), salt (grey) and in RLN ratio (salt/control; orange) for tested accessions listed in Table S1. C. Manhattan plots for flowering time ratio (salt/control), salt and control conditions (See Table S2). D. Manhattan plots for RLN ratio (salt/control), salt and control conditions (See Table S2).

**Figure S2.**
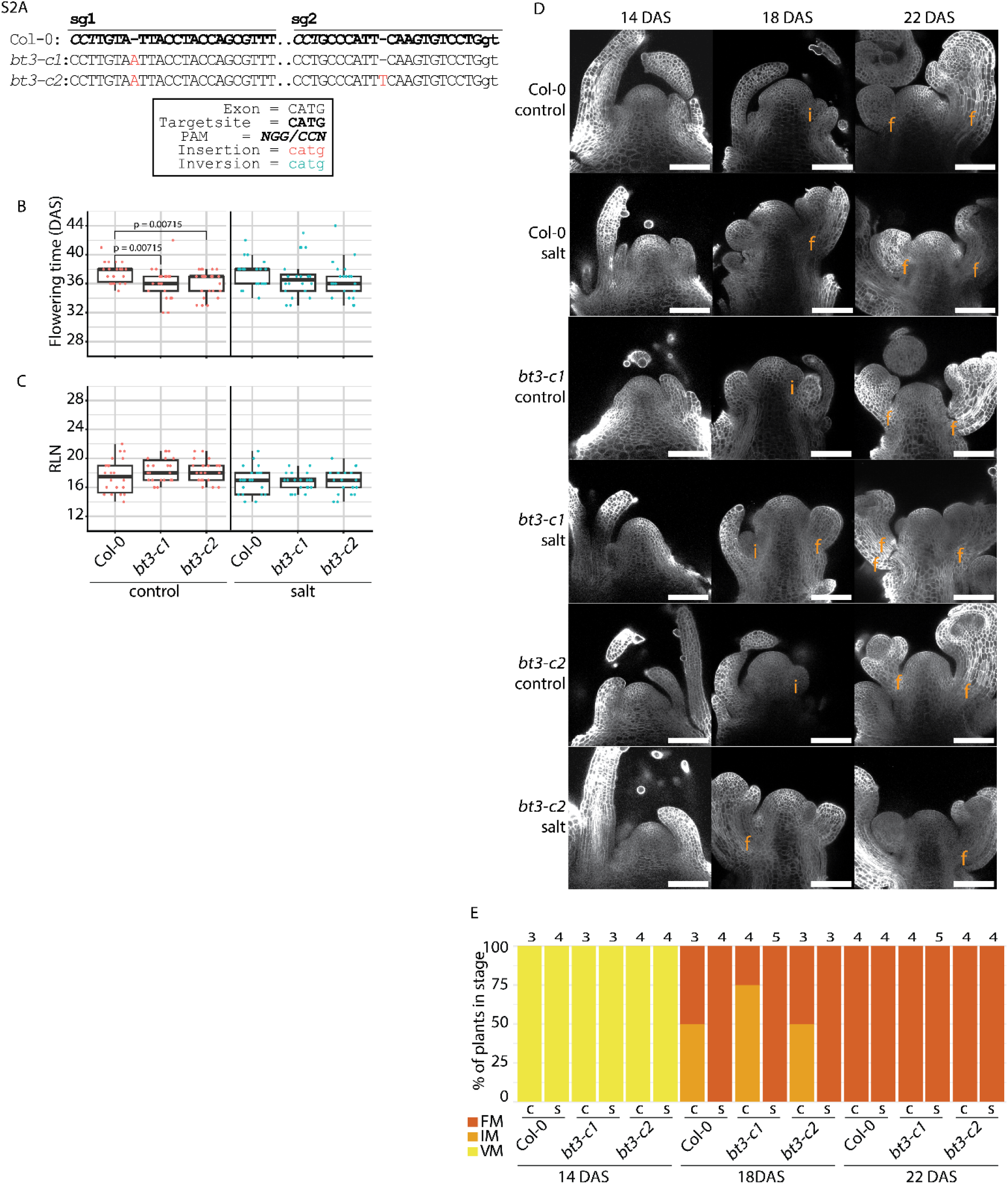
The floral transition in *bt3* mutants A. CRISPR-mutations in *bt3-c1 & -c2*. B & C. Flowering time and RLN for genotypes grown climate-controlled growth chambers in LD in control (red) and salt (blue) conditions (Data are pooled from two independent experiments, n = 20 – 23). P-values within control or salt treatment indicate significant differences between genotypes using Pairwise Wilcoxon Rank Sum Test with Benjamini-Hochberg multiple testing correction. Data corresponds to experiments shown in Figure 2B&C. D. Representative confocal images of the SAMs of the respective genotypes grown in climate-controlled growth chambers in LD during the transition from a vegetative meristem to a reproductive meristem. Inflorescence meristems and floral meristems indicated with “i” and “f”, respectively. Scale bar = 100 µm. E. Quantification of SAM developmental stages from vegetative meristem (VM) to an inflorescence meristem (IM) and floral meristem (FM) in control (c) an salt (s) conditions. Numbers above bars indicate the total number of replicates per genotype/treatment/timepoint.

**Figure S3.**
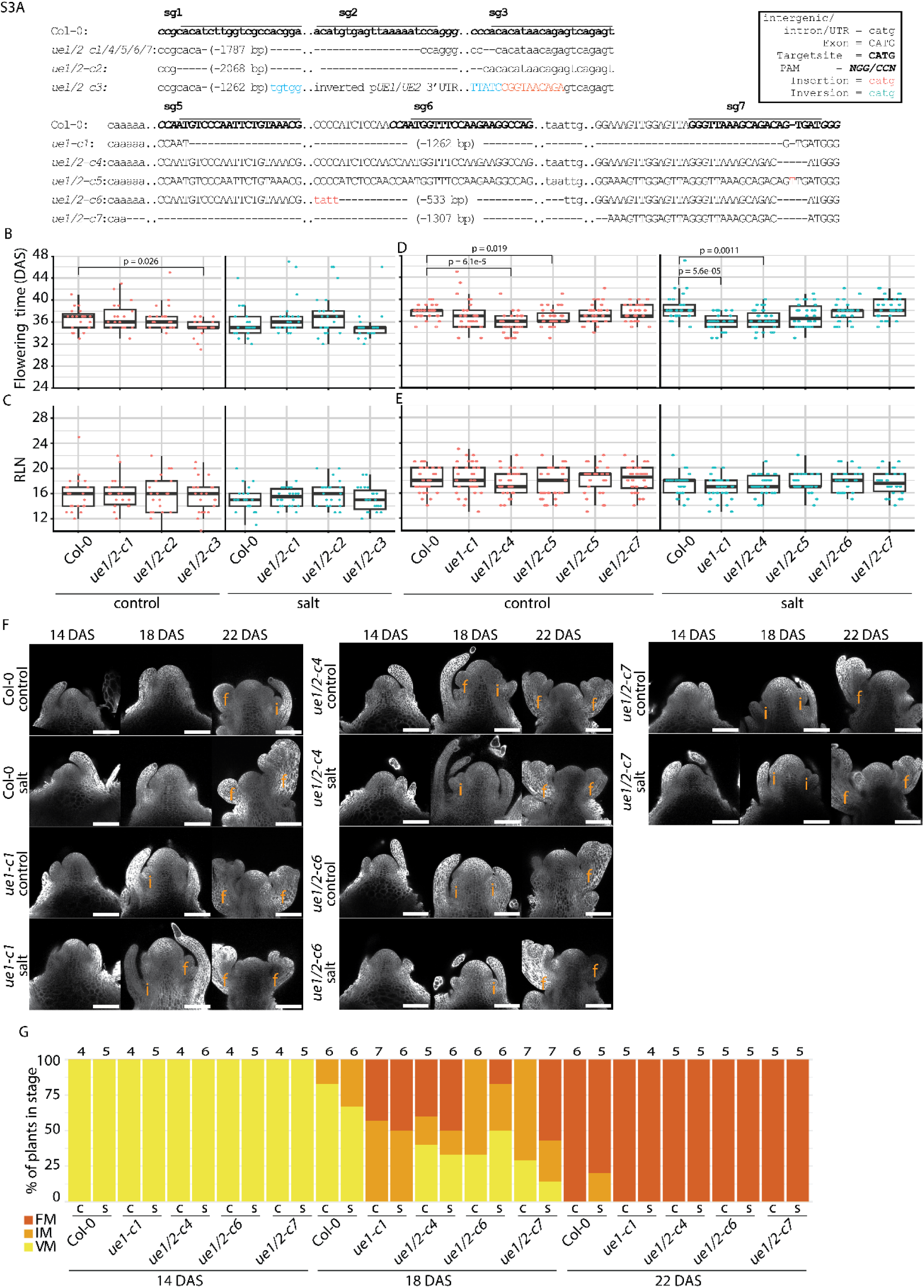
*UGT74E1* and *UGT74E2* delays bolting time in control conditions A. CRISPR-mutations in *ue1-c1 and ue1/2-c1 to ue1/2-c7*. B-E. Flowering time and RLN for genotypes grown in climate-controlled growth chambers in LD in control and salt conditions (Data are pooled from two independent experiments, B&C: n = 18 – 23; D&E: n = 25 - 31). P-values within control or salt treatment indicate significant differences between genotypes using Pairwise Wilcoxon Rank Sum Test with Benjamini-Hochberg multiple testing correction. Data corresponds to experiments shown in Figure 3D-F. F. Representative confocal images of the SAMs of the respective genotypes grown in in climate-controlled growth chambers in LD during the transition from a vegetative meristem to a reproductive meristem. Inflorescence meristems and floral meristems indicated with “i” and “f”, respectively. Scale bar = 100 µm. G. Quantification of SAM developmental stages from vegetative meristem (VM) to an inflorescence meristem (IM) and floral meristem (FM) in control (c) and salt (s) conditions at indicted timepoints. Numbers above bars indicate the total number of replicates per genotype/treatment/timepoint.

**Figure S4.**
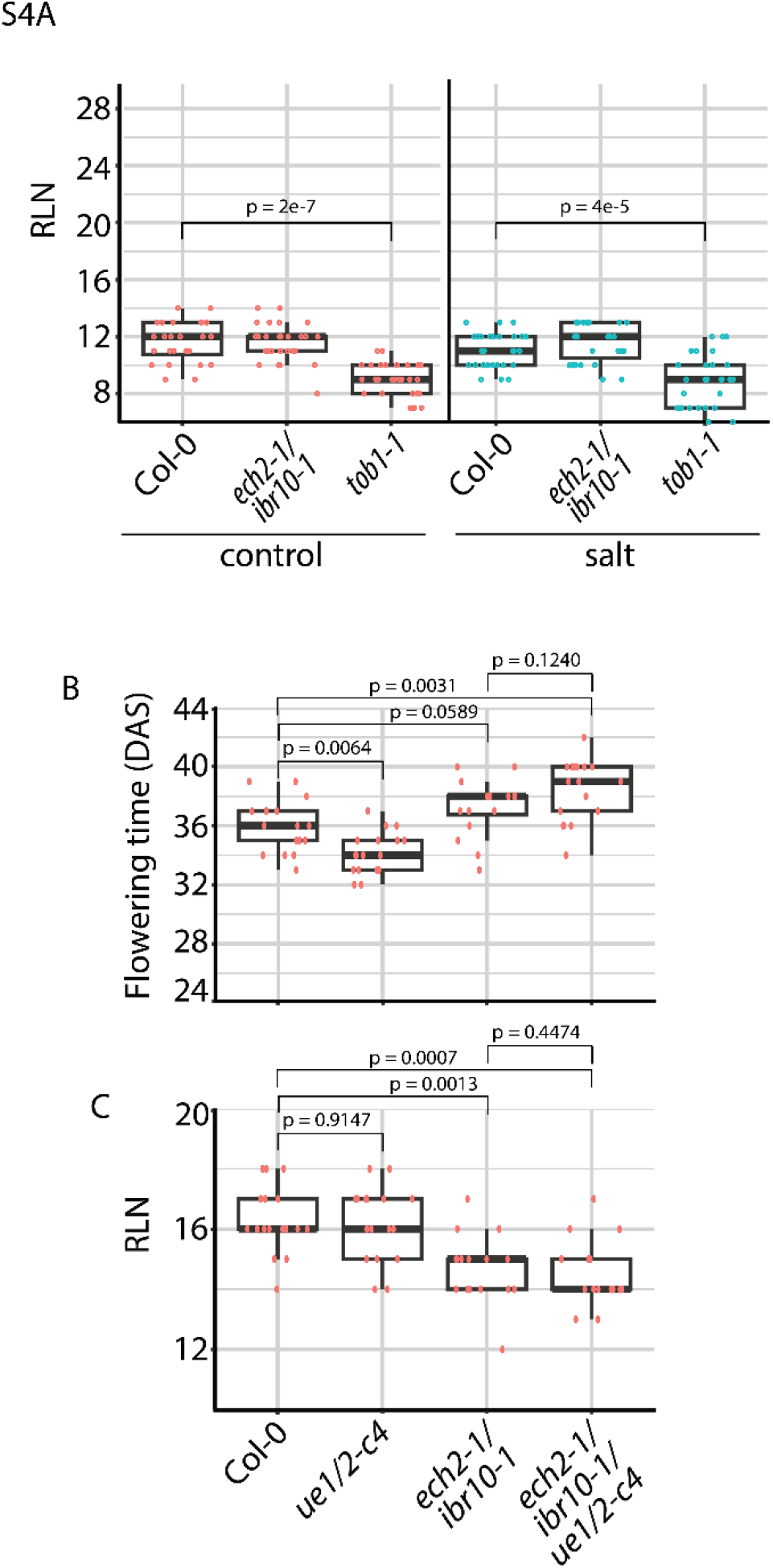
Bolting and flowering time regulation by *UGT74E1 & UGT74E2* requires IBA catabolic enzymes ECH2/IBR10 A RLN for genotypes grown in a climate-controlled greenhouse with supplemented light in LD in control (red) and salt (blue) conditions (Data are pooled from two independent experiments, n = 27 – 28). P-values within control or salt treatment indicate significant differences between genotypes using Pairwise Wilcoxon Rank Sum Test with Benjamini-Hochberg multiple testing correction. Data corresponds to experiments shown in Figure 4A-C. B&C. Flowering time and RLN for genotypes grown in climate-controlled growth chambers in LD in control conditions (Data from one independent experiment, n = 16 – 17). P-values indicate statistical differences between genotypes using Pairwise Wilcoxon Rank Sum Test with Benjamini-Hochberg multiple testing correction. Data corresponds to experiments shown in Figure 4D&E.

**Figure S5.**
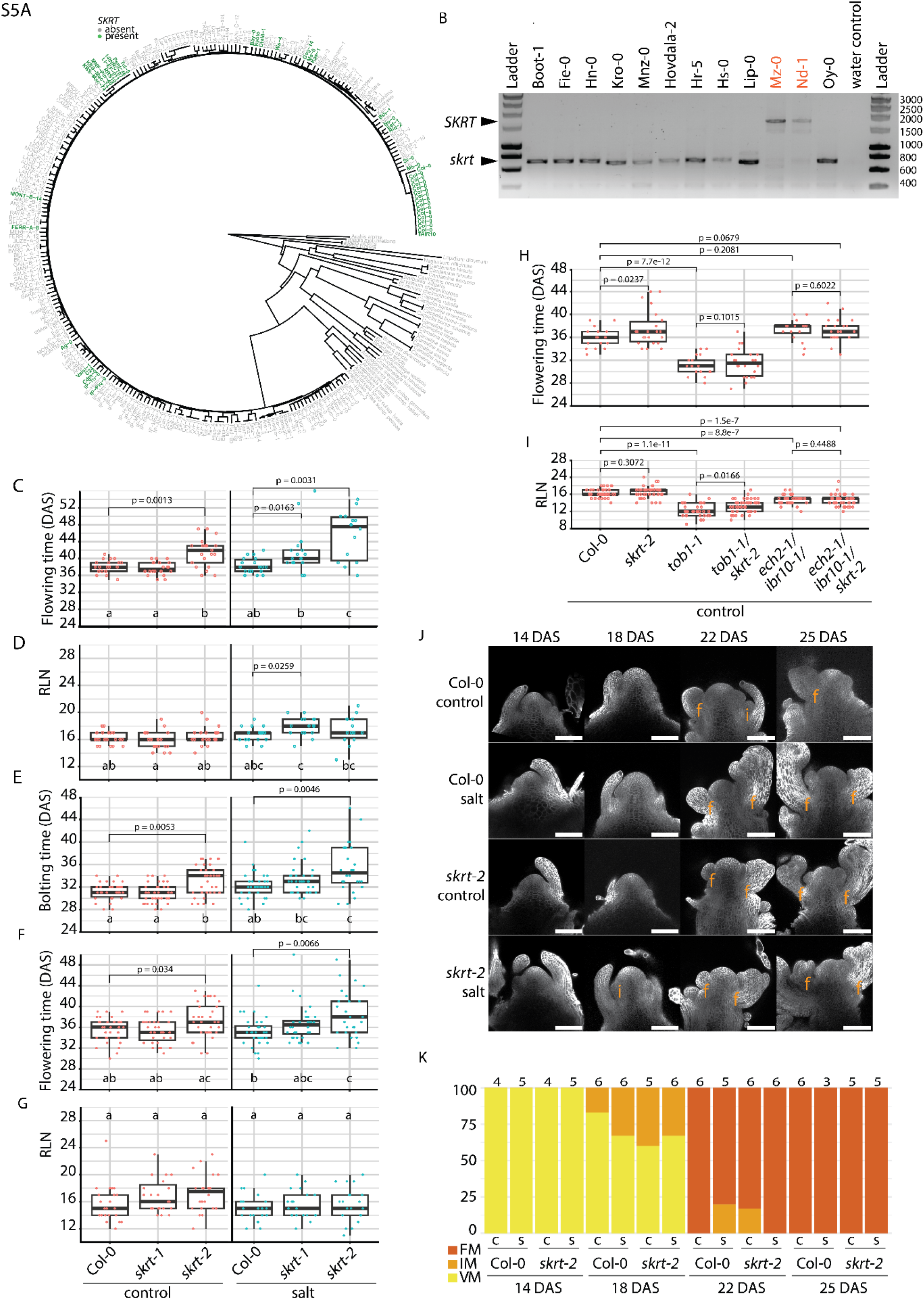
The DNA transposon *SAUERKRAUT (SKRT)* accelerates bolting time in control and salt. A. Phylogenetic analysis of *SKRT* in 287 genomes of *Arabidopsis thaliana* and related species. Minimum Evolution tree based on whole-genome MinHash distances between all input genomes. Accessions in green contain the *SKRT* insertion and grey accessions do not. See Table S4 for a larger version of this tree and more information on the represented genotypes. B. PCR on *SKRT* using JAD010-JAD050 (see Materials and Methods; Table S15) on gDNA from 12 Arabidopsis accessions. In black, accessions showing the highest bolting time ratio (salt/control) that carry the B allele (Figure 1E, Table S1) all lack *SKRT* (∼ 750 bp fragment). In red, Mz-0 and Nd-1 that are not delayed in bolting time ratio (salt/control) and contain the A allele (Figure 1E, Table S1) do contain *SKRT* ( ∼ 1750 bp fragment). C&D. Flowering time and RLN for genotypes grown in LD in control (red) and salt (blue) conditions (Data from one independent experiments, n = 14-21, Figure S5 shows more independent replicates). P-values within control or salt treatment indicate significant differences between genotypes using Pairwise Wilcoxon Rank Sum Test with Benjamini-Hochberg multiple testing correction. Letters indicate significant (p < 0.05) differences between genotypes*treatments using a two-way ANOVA using Tukey-HSD multiple testing correction. Data corresponds to experiments shown in Figure 5D&E. E-G. Independent experimental replicates for *skrt-2* mutants phenotypes: bolting time, flowering time and RLN. Genotypes were grown in climate-controlled growth chambers in LD in control (red) and salt (blue) conditions. E&F: Data are pooled from three independent experiments, n = 28 – 35. G: Data are pooled from two independent experiments, n = 20 -23. P-values within control or salt treatment indicate significant differences between genotypes using Pairwise Wilcoxon Rank Sum Test with Benjamini-Hochberg multiple testing correction. Letters indicate significant (p < 0.05) differences between genotypes*treatments using a two-way ANOVA using Tukey-HSD multiple testing correction. H&I. Flowering time and RLN for genotypes grown in LD in climate-controlled growth chambers at control conditions (Data from two independent experiment, n = 33-43). P-values indicate statistical differences relative to control genotypes as determined by Pairwise Wilcoxon Rank Sum Test with Benjamini-Hochberg multiple testing correction. Data corresponds to experiments shown in Figure 5F&G. J. Representative confocal images of the SAMs of the respective genotypes during the transition from a vegetative meristem to a reproductive meristem. Inflorescence meristems and floral meristems indicated with “i” and “f”, respectively. Scale bar = 100 µm. K. Quantification of SAM developmental stages from vegetative meristem (VM) to an inflorescence meristem (IM) and floral meristem (FM) in control (c) an salt (s) conditions. Numbers above bars indicate the total number of replicates per genotype/treatment/timepoint.

**Figure S6.**
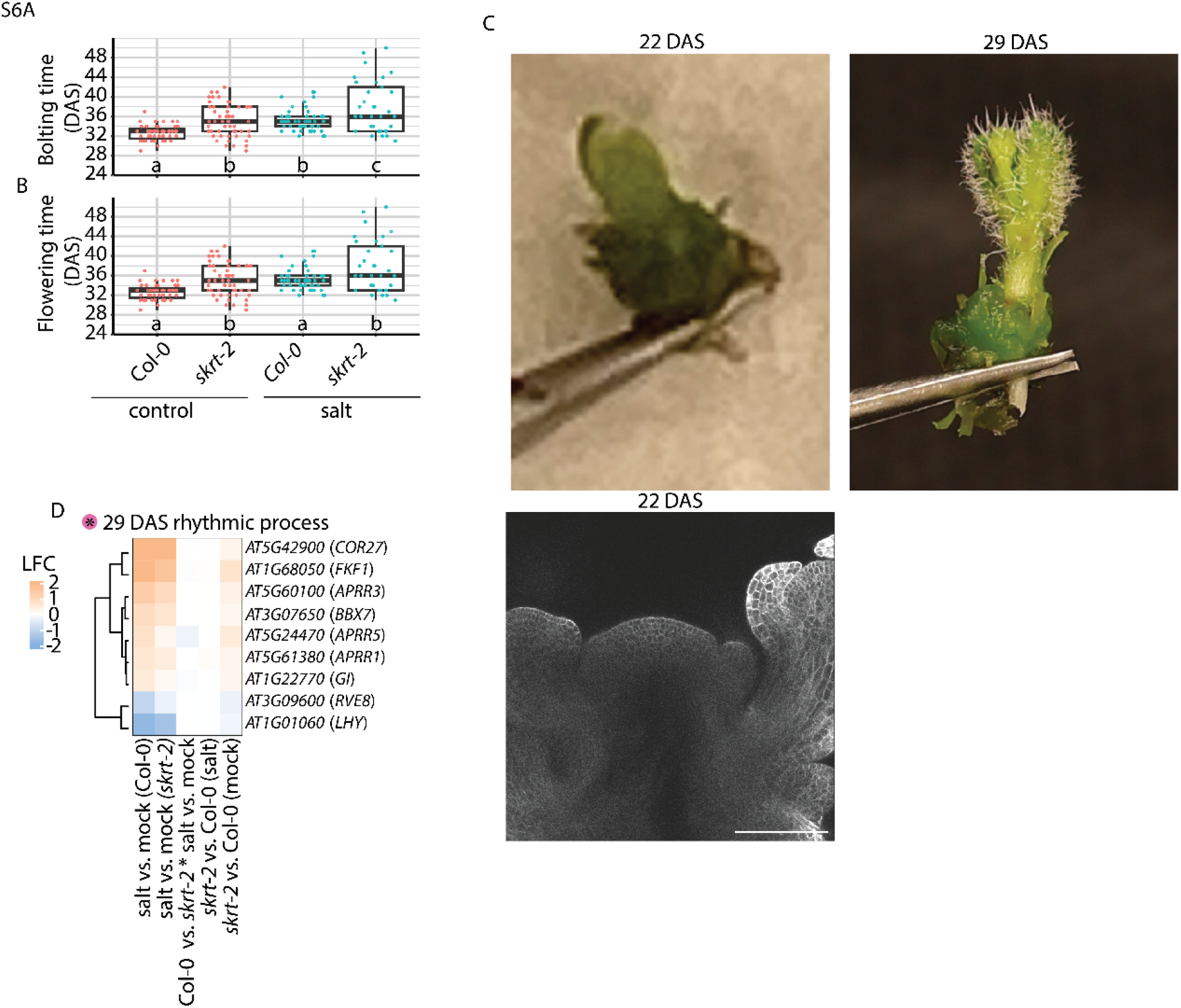
Harvested tissue, developmental stages and floral transition phenotypes of RNA-seq samples A & B. Bolting and flowering time data (n= 33-52) for Col-0 and *skrt-2* grown together with plants that were used for RNA-seq. Letters indicate significant (p < 0.05) differences between genotypes*treatments using a two-way ANOVA using Tukey-HSD multiple testing correction. C. Top: Representative images of harvested shoot apices for RNA-seq for 22 DAS and 29 DAS, and bottom: representative confocal microscopy image (Scale bar = 100 µm) of an induced SAM with floral meristems at 22 DAS. All images were harvested from Col-0 and *skrt-2* plants grown together with plants that were used for RNA-seq. D. Heatmap showing the relative genes expression as log_2_ fold change (LFC) for five contrasts of genes listed in 29 DAS GO:BP “rhythmic process” (See pink dot with asterisk in figure 6B). See Table S6 for LFC values and p-adjusted values for all DEGs in all contrasts.

**Figure S7.**
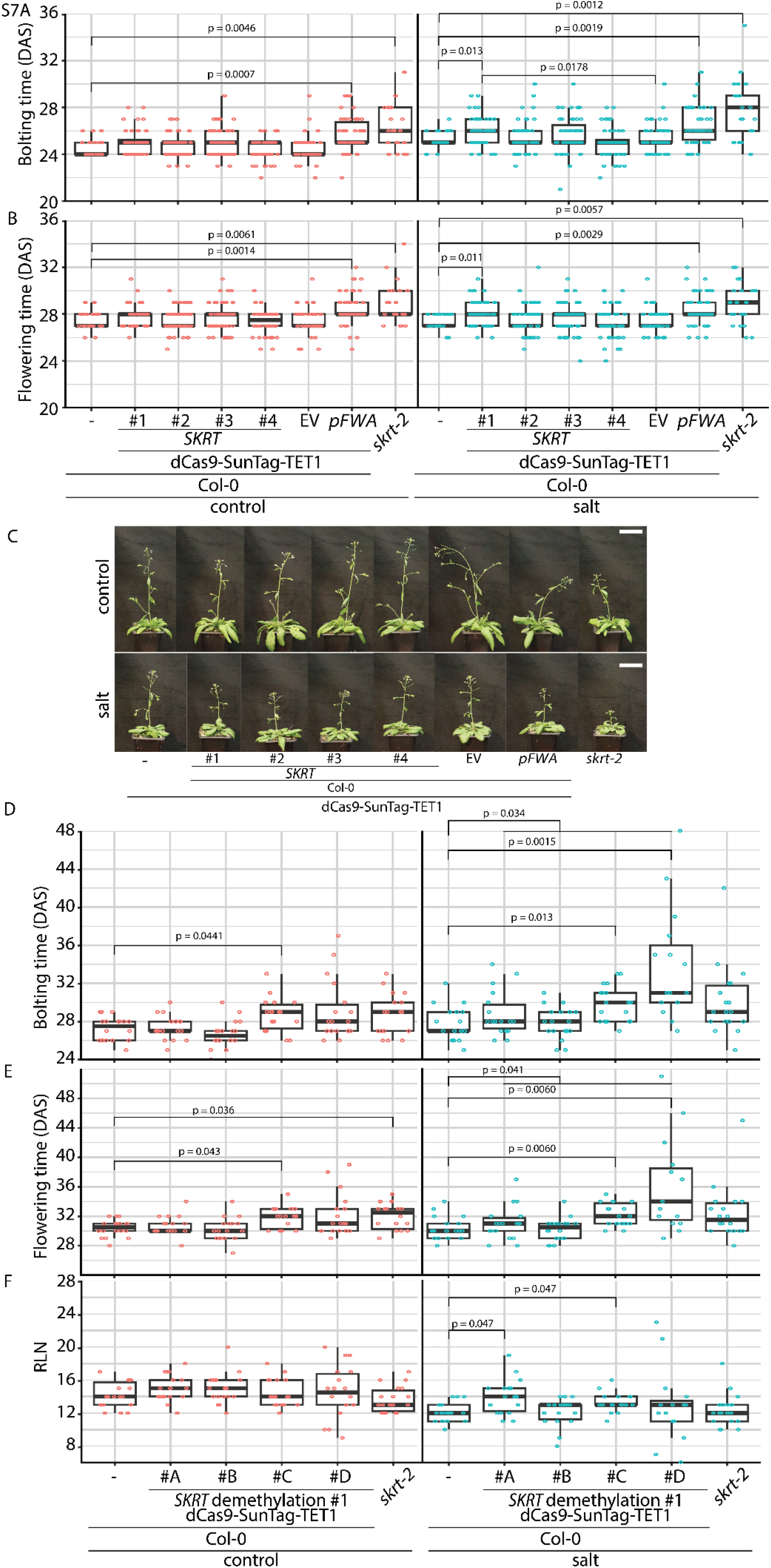
Demethylation of *SKRT* leads to a salt-dependent delay in bolting time. A-B. Bolting time and flowering time for genotypes grown in a climate-controlled greenhouse with supplemented light in LD in control (red) and salt (blue) conditions (matches with Figure 7C). For Col-0 and *skrt-2* 21 replicates per genotype per treatment were used, for *dCas9-SunTag-TET1* lines 20 independent T2 families with 2-3 replicates per family were used per genotype per treatment (n = 44 - 59). P-values within control or salt treatment indicate significant differences between genotypes using Pairwise Wilcoxon Rank Sum Test with Benjamini-Hochberg multiple testing correction. Data corresponds to experiment shown in Figure 7C&D. C. Representative image of shoot architecture at 31 DAS of genotypes shown in A&B, grown in a climate-controlled greenhouse with supplemented light in LD in control and salt conditions. D-F. Bolting & flowering time and RLN for genotypes grown in a climate-controlled greenhouse with supplemented light in LD in control (red) and salt (blue) conditions. For Col-0 and *skrt-2* 21 replicates per genotype per treatment were used, for *dCas9-SunTag-TET1* lines 4 independent T2 families with 15-18 replicates per family were used per genotype per treatment. P-values within control or salt treatment indicate significant differences between genotypes using Pairwise Wilcoxon Rank Sum Test with Benjamini-Hochberg multiple testing correction.

## Notes

### Competing Interest Statement

The authors have declared no competing interest.

